# Tunable Bessel beam two-photon fluorescence microscopy for high-speed volumetric imaging of brain dynamics

**DOI:** 10.64898/2025.12.31.697185

**Authors:** Mengyang Jacky Li, Jinghui Wang, Mikolaj Walczak, Yuqing Qiu, Colleen Russell, Miroslaw Janowski, Piotr Walczak, Yajie Liang, Tian-Ming Fu

## Abstract

High-speed volumetric imaging of the brain is essential for linking diverse cellular events to tissue-level functions. However, the brain’s structural and dynamic heterogeneity—spanning microns to millimeters and milliseconds to hours—requires imaging techniques with tunable spatiotemporal resolution, flexible 3D sampling, and compatibility with targeted perturbations. Here, we present tunable Bessel beam two-photon fluorescence microscopy (tBessel-TPFM), a compact, low-cost, and versatile platform for intravital brain imaging across millimeter scale with subcellular resolution. tBessel-TPFM transforms slow 3D volume scans into fast 2D frame scans via an axially elongated Bessel focus, achieving acquisition rates ∼100-fold faster and reduced motion artifacts compared with conventional TPFM. Exploiting its full tunability of the Bessel focus, we applied tBessel-TPFM for quantitative mapping of cerebral blood flow and neurovascular coupling in normal and ischemic stroke mice. Unlike existing Bessel focus generation methods, the axial center of tBessel-TPFM remains fixed at the objective focal plane during profile tuning. Leveraging this advantage, we integrated tBessel-TPFM with simultaneous 3D targeted optogenetic stimulation for volumetric neuronal connectivity mapping. We also tracked microglial process dynamics following single-cell laser ablation, revealing diverse neuroimmune responses across spatial and temporal scales. By combining high speed, deep penetration, tunable sampling, and multimodal perturbation, tBessel-TPFM empowers a broad spectrum of neurobiological investigations—from vascular physiology and functional connectivity to neuroimmune interactions.

## Introduction

The brain’s normal function relies on a dynamic interplay between diverse biological processes. These interactions span across spatial and temporal scales, from blood flows within hierarchical vascular networks and communication among neurons forming complex circuits to the constant surveillance for damage and pathogens by glial cells. Studying these processes requires the ability to observe volumetric dynamics deep within highly scattering brain tissues. Moreover, each process imposes its distinct imaging demands. For example, hemodynamics mapping requires only moderate spatial resolution to visualize capillaries but demands acquisition rates up to the kilohertz range to capture fast blood flows in arteries and veins. In contrast, imaging microglial dynamics requires subcellular resolution to identify fine processes, but only sub-second temporal resolution. A versatile, cost-effective imaging platform capable of capturing volumetric biological dynamics in the brain while accommodating such diverse requirements could facilitate discoveries across neuroscience, neurology, brain immunology, and oncology. Furthermore, integrating spatiotemporally patterned perturbations into such a platform would enable simultaneous targeted interventions and volumetric investigations, providing broad-reaching implications for the development of novel therapeutic strategies.

In recent years, several promising brain imaging technologies have emerged. Functional magnetic resonance imaging offers whole-brain coverage, but lacks cellular resolution(Aries et al., 2010; D’Esposito et al., 2003). Optical coherence tomography allows label-free contrast with moderate penetration depth, yet falls short to resolve subcellular structures(Drexler et al., 2001; Tang et al., 2025). Time-resolved laser speckle imaging, while offering better spatial resolution, suffers from limited depth(Vaz et al., 2016). Photoacoustic imaging can penetrate deeper, but its depth-dependent resolution complicates data interpretation(Lin and Wang, 2022).

Two-photon fluorescence microscopy (TPFM), which scans a spatially confined Gaussian focus across a three-dimensional (3D) volume, can achieve subcellular resolution deep within highly scattering brain tissues(Shih et al., 2012; Scheele et al., 2022; Xu et al., 2024). As such, TPFM has been widely adopted for *in vivo* brain imaging, revealing insights from synaptic plasticity to glia-neuron interactions(Scheele et al., 2022; Xu et al., 2024). Nevertheless, scanning the focus across all three dimensions results in slow volumetric acquisitions(Wu et al., 2021). To address this, strategies such as remote focusing, spatiotemporal multiplexing, and projection imaging have been developed(Kim and Schnitzer, 2022; Weisenburger and Vaziri, 2018; Wu et al., 2021). Notably, an elegant approach—Bessel beam TPFM—employs an axially elongated Bessel focus and converts a 3D volumetric scan into a 2D frame scan, enabling video-rate volumetric imaging with diffraction-limited resolution(Lu et al., 2017). Despite its advantages, current Bessel beam generation methods limit the widespread adoption of Bessel beam TPFM in biomedical research. Systems based on a single axicon-lens pair lack the tunability to meet diverse brain imaging demands. Spatial light modulator (SLM)-based approaches offer tunability, but are prohibitively expensive, incompatible with simultaneous multi-wavelength excitation, and suffer from significant light loss. Several methods exploiting the combinations of axicons and lenses have been reported(Lu et al., 2018; Schnieder et al., 2024; Takanezawa et al., 2021), but they all introduce axial shifts of the Bessel beam during beam profile tuning. These unwanted focal shifts necessitate frequent system realignment and prevents integration with simultaneous optogenetic stimulation or laser ablation(Wang et al., 2016).

Here, we overcome these limitations by introducing tunable Bessel beam two-photon fluorescence microscopy (tBessel-TPFM), a compact, low-cost, light-efficient, and versatile platform for intravital imaging of diverse brain dynamics across millimeter scale with subcellular resolution. Importantly, tBessel-TPFM allows for adaptive tuning of the Bessel beam’s spatial resolution, volume coverage, and energy confinement to optimize for specific imaging needs. Leveraging this tunability, we quantitatively measured cerebral blood flows in normal and ischemic stroke mice across spatial and temporal scales—from slow blood flows in tiny capillaries to ultrafast flows in arteries and veins. With implemented focal jumping and lateral tiling, we monitored capillary blood flow and diameter changes in real time across a 2,500×2,500×450-μm^3^ cortical volume, and imaged neurovascular coupling (NVC) across cortical layers in awake mice. Furthermore, unlike existing methods, tBessel-TPFM maintains the axial center of the Bessel beam at the objective focal plane during profile tuning, enabling integration with 3D targeted optogenetic stimulation for volumetric neuronal connectivity mapping. Finally, by combining high spatial resolution, rapid volumetric acquisition, and simultaneous two-photon perturbation, we tracked microglial processes dynamics following single-cell laser ablation, revealing rapid neuroimmune responses.

## Results

### Tunable Bessel beam two-photon fluorescence microscopy

In contrast to conventional Gaussian TPFM that scans an axially confined Gaussian focus across a 3D volume (Figure 1A, top), Bessel-TPFM images the same volume by scanning an axially extended Bessel beam across a 2D plane (Figure 1A, bottom)(Lu et al., 2017). When the Bessel beam length matches the depth of the imaging volume, the resulting image corresponds to a 2D projection of the 3D stack that would otherwise require 10–100 sequential z-slices using conventional Gaussian TPFM, therefore achieving a 10–100× increase in acquisition speed by capturing the entire volume in a single frame(Fan et al., 2020). For hemodynamics and functional imaging where the biological structures—vasculatures and neurons—remain largely unchanged, a two-step strategy is adopted: first acquire a high-resolution 3D structural reference with conventional Gaussian TPFM, then perform dynamic volumetric imaging with Bessel-TPFM (Figure 1A). The three core imaging metrics of Bessel-TPFM—spatial resolution, temporal resolution, and structural overlap—are inherently linked in a triangular trade-off (Figure 1B). Optimizing these competing metrics for different applications, such as capillary flow versus neuronal activities, requires the ability to adjust the Bessel beam properties independently.

**Figure 1:**
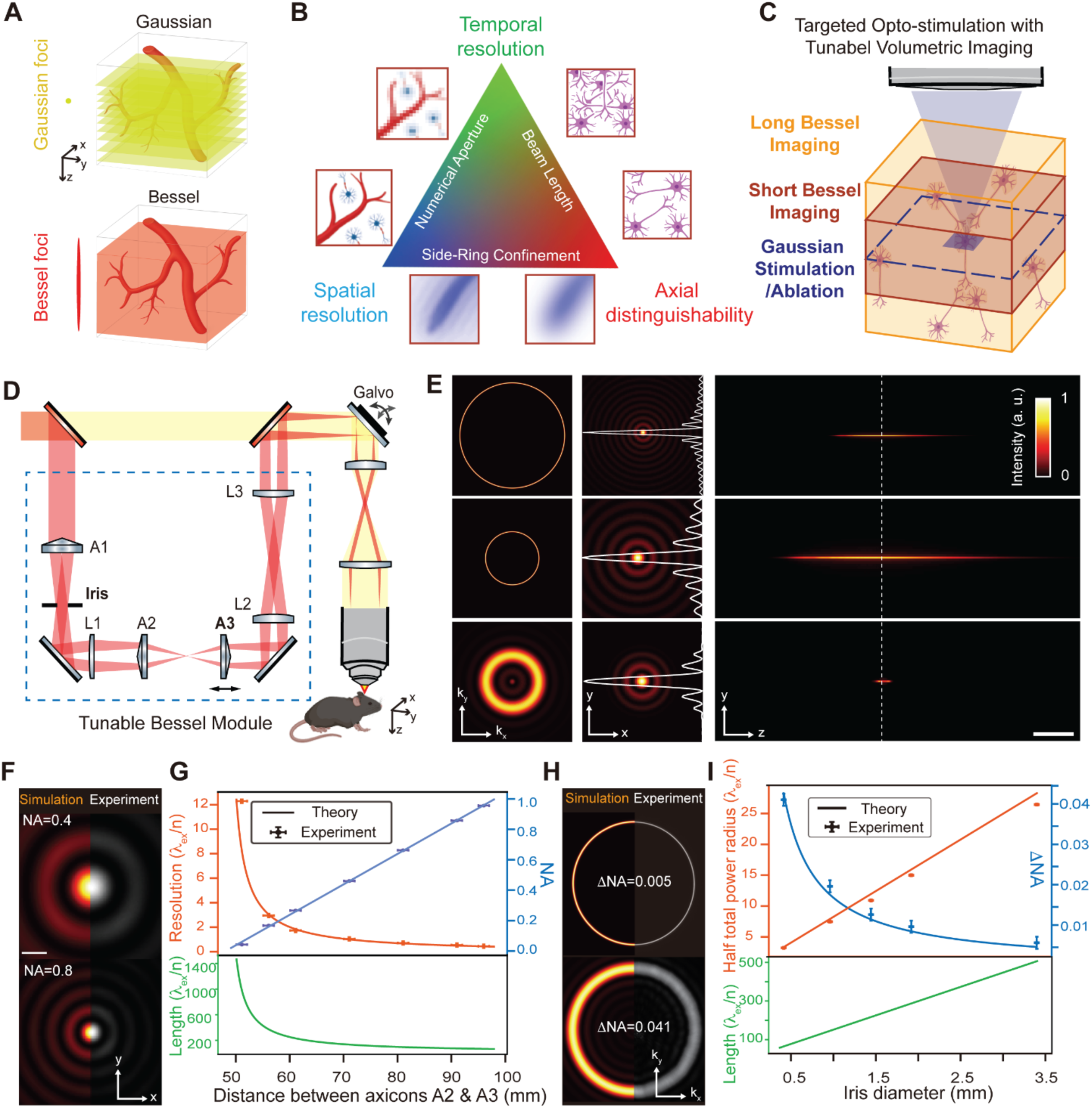
Concept, design, and characterization of tunable Bessel two-photon microscopy. (**A**) Comparison of Gaussian (top) and Bessel (bottom) two-photon fluorescence microscopy (TPFM). Gaussian imaging requires sequential acquisition of 2D images at different axial positions, whereas Bessel imaging captures the entire volume in a single frame. (**B**) Fundamental trade-offs in Bessel beam imaging between spatial resolution, temporal resolution, and axial distinguishability, governed by the numerical aperture (NA), beam length, and side-ring confinement. (**C**) For targeted opto-stimulation, Bessel beam tuning must preserve a fixed axial beam center to maintain co-registration with stimulation or ablation beams. (**D**) Schematic of the tunable Bessel module (A1–A3, axicons; L1–L3, lenses; iris) and its integration with Gaussian-TPFM. A1 with L1 generates the Bessel beam; the iris adjusts ΔNA; A2–A3 spacing sets NA; L2–L3 form a 4f relay to the scanning galvo. (**E**) Zemax simulations showing pupil (left), sample (middle), and propagation (right) images for 0.8 NA with open iris (top), 0.4 NA with open iris (middle), and 0.4 NA with 10% iris opening (bottom). (**F**) Simulated versus experimental Bessel beam cross-section profiles at NA= 0.4 (top) and 0.8 (bottom). Experimental images were acquired using a camera conjugate to the objective’s sample plane. Scale bar: λ/n, where n is the refractive index of the immersion medium. (**G**) Theoretical and experimental results showing adjustment of the Bessel beam NA (blue), resolution (orange), and length (green) as a function of A2-A3 spacing. (**H**) Simulated versus experimental Bessel beam pupil profiles for ΔNA=0.005 (top) and 0.041 (bottom). Experimental images were acquired using a camera conjugate to the objective’s back focal plane. (**I**) Theoretical and experimental results showing adjustment of the Bessel beam ΔNA (blue), half total power radius (orange), and length (green).

In practice, these metrics are governed by three key parameters of the Bessel beam: lateral resolution, axial beam length, and side-ring confinement. First, lateral resolution, set by the numerical aperture (NA) of the Bessel beam, determines the ability to resolve fine structures but comes at the cost of speed, as higher resolution requires smaller scanning step size (Figure 1B, left). Optimal spatial resolution is therefore task-dependent: fine structural dynamics benefit from higher resolution, whereas ultrafast flow measurements favors faster frame rates. Second, axial beam length, set by the NA and NA broadening (ΔNA) of the beam, governs the depth coverage of the imaging volume, but at the cost of structural overlap: Longer beams expand depth coverage at the same acquisition speed but can confound signals from closely spaced structures. (Figure 1B, right). Third, Bessel beam side-ring add another constraint: high-NA, long Bessel beams generate substantial side-ring excitation, leading to potential imaging artifacts (Figure 1B, bottom)(Chen et al., 2024).

While several elegant methods that can tune lateral resolution, axial beam length, and side-ring confinement exist, they all suffer from a key limitation: adjusting the Bessel beam profile shifts the beam axially, displacing the imaging volume (Figure S1 in supplementary file 1)(Dickey and Conner, 2011; Lu et al., 2018; Takanezawa et al., 2021). Although this undesired displacement is acceptable to certain imaging applications, it forces realignment, complicates experiments, and reduces reproducibility. Moreover, it is particularly problematic when imaging is combined with optogenetic stimulation or laser ablation, where precise co-registration of stimulation and imaging volumes is required (Figure 1C). Minimizing axial shift while tuning the Bessel beam profile is therefore a core requirement to enable volumetric perturbation–imaging studies.

To meet this requirement, we designed a compact, tunable Bessel (tBessel) module that can be readily integrated into conventional Gaussian TPFM (Figure 1D). For broad adoptability and high light efficiency, the tBessel module uses only off-the-shelf components, avoiding programmable or diffractive elements (Figure S2 in supplementary file 1). The optical design consists of three major parts. First, an axicon-lens pair (A1 and L1) converts a collimated beam into a focused ring beam—Fourier transform of a Bessel beam—at a rear pupil-conjugate plane. Second, to enable tunability, the diameter and thickness of the ring beam—corresponding to the NA and ΔNA, respectively—needs be adjustable. Our tBessel module achieves NA tuning by exploiting a pair of axicons (A2 and A3, Figure 1D), where the separation between the two controls the NA. Importantly, varying this separation introduces minimum phase variation (Figure S3, and supplementary note 2 in supplementary file 1), resulting in negligible axial shift (Figure 1E; Figure S2C, video S1, and Supplementary Note 1 in supplementary file 1). To enable independent adjustment of ΔNA, an iris is conjugated to the objective focal plane (Figure 1E; Figure S4, and video S1 in supplementary file 1). This decoupled control of NA and ΔNA allows independent tuning of the Bessel beam’s spatial resolution and side-ring confinement (Figure 1E and video S1 in supplementary file 1).

Next, we characterized the tBessel module experimentally. First, by varying the axicon pair separation, the resolution of the Bessel beam can be adjusted in close agreement with numerical simulations (Figure 1F). Quantitatively, spatial resolution scales inversely with the axicon pair separation, while axial beam length varied accordingly (Figure 1G). Second, changing the iris aperture alters the ΔNA, as evidenced by the pupil profile images that match numerical simulations (Figure 1H). Quantitatively, ΔNA scales inversely with iris aperture, and both axial beam length and side-ring confinement radius—defined as the radius where half of the total power is contained—vary linearly with the iris aperture (Figure 1I and Figure S5 in supplementary file 1). Noticeably, across the full tuning range, experimental and simulated results showed excellent agreement in lateral resolution, axial beam length, and side-ring confinement. Finally, the tBessel module is capable of simultaneous multi-wavelength excitation—an advantage over SLM-based methods (Chen et al., 2014; Lu et al., 2017). As a demonstration, we generated 520-nm and 635-nm Bessel beams concurrently and observed negligible chromatic shift over a broad NA range (Figure S2, E to H in supplementary file 1). This minimum chromatic aberration minimizes pulsed laser dispersion, enhancing fluorescence excitation efficiency.

### Cortical-wide imaging of vascular dynamics and neurovascular coupling

tBessel-TPFM enables rapid volumetric imaging, but its axial beam length must be optimized. Overly long beams cause structural overlap that obscures signal localization, while overly short beams underutilize their volumetric speedup (Figure 2A). To determine the optimal beam length for vascular imaging, we quantified axial inter-vessel distances (AIV) at different cortical depths from 3D Gaussian-TPFM data. Gamma-function fits revealed depth-dependent AIV (Figure S6 in supplementary file 1). This highlights the need for flexible tuning of axial beam length. To cover extended regions, we combined tBessel-TPFM with lateral tiling and piezo-driven axial jumps. Using a 0.4 NA, 150-μm-long Bessel beam, each scan covered 700×700×150-μm^3^ in 33-ms. With 4×4 lateral tiles and three axial jumps, we stitched 48 volumes spanning 2,500×2,500×450-μm^3^ (Figure 2B). Depth-color-coded reconstructions revealed large superficial vessels (0–150-μm) and dense capillary networks at deeper layers (150–450-μm) with penetrating arterioles and venules clearly resolved (Figure 2C). Time-lapse imaging enabled visualization of vasodilation and vasoconstriction dynamics (Figure 2C and video S2 in supplementary file 1), allowing further investigation of neurovascular coupling (NVC).

**Figure 2:**
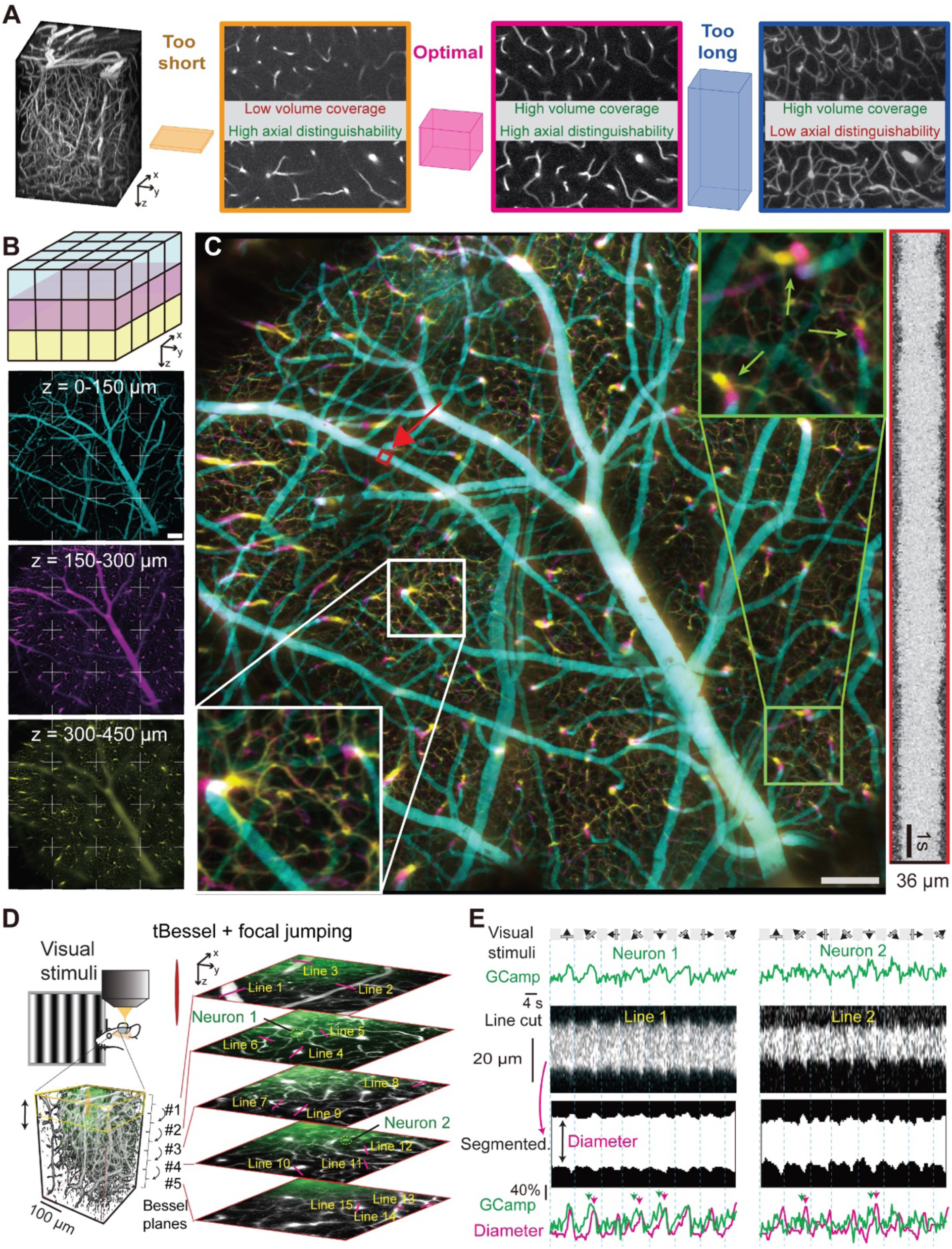
Large-scale volumetric vascular imaging and neurovascular coupling. (**A**) Left: 3D rendering of a 350×350×450-μm^3^ Gaussian stack showing dense cortical vasculature. Right: representative Bessel images with beam lengths of 17-μm (orange), 35-μm (magenta), and 140-μm (blue), illustrating the trade-off between information content and structural overlap. (**B**) tBessel scan combining lateral tiling (4×4) and focal jumps (3, color-coded), yielding 48 tiles covering a 2,500×2,500×450-μm^3^ cortical volume. Scale bar: 200-μm. (**C**) Combined millimeter-scale tBessel image from (**B**). White box: sub-capillary resolution. Green box: penetrating arterioles/venules (arrows). Right: vasoconstriction and vasodilation of the vessel marked in red. Scale bar: 200-μm. (**D**) Experimental setup for functional imaging of primary visual cortex (V1). Mice with cranial windows over V1 were presented moving gratings in eight directions. Five planes spanning a 260×260×400-μm^3^ volume were acquired at 2.5-Hz (5 trials per direction). Two neurons were analyzed for fluorescence dynamics, and vessels were outlined for resliced space–time plots of diameter changes. (**E**) Top: averaged calcium traces (green) from two neurons. Middle: raw and segmented resliced images from lines 1 and 2 in (**D**). Bottom: overlay of neuronal activity (green) and vessel diameter (magenta) showing strong correlation, with vascular changes lagging neuronal responses (arrows).

We investigated NVC in the primary visual cortex (V1) of awake mice using tBessel-TPFM with focal jumping (Figure 2D). NVC links neuronal activation to rapid increases in blood flow that supply nutrients and clear metabolites, and its disruption is implicated in neurological and psychiatric disease(Iadecola, 2004; Kaplan et al., 2020; Kisler et al., 2017; Schaeffer and Iadecola, 2021). We simultaneously monitored neuronal activity with GCaMP and vascular dynamics across cortical layers during visual stimulation(Niell and Stryker, 2008) (Figure 2D and Figure S7A). From Gaussian reference stacks (260×260×400-μm^3^), we determined an 80-μm long Bessel beam with five focal jumps as optimal, balancing imaging speed and axial overlap (Figure S7 B and C in supplementary file 1). With this configuration, we imaged neuronal calcium and vascular dynamics at 3-Hz while presenting moving gratings to the mice. Trial-averaged calcium traces showed strong direction selectivity (Figure 2E, top). To assess vascular responses, 15 vessels spanning arterioles, venules, and capillaries were analyzed across imaging planes (Figure 2D). Space–time plots from line-cuts revealed clear dilation in arterioles but little change in neighboring venules (Figure 2E, middle; Figure S7 E and F in supplementary file 1). Vasodilation consistently lagged neuronal activity (Figure 2E, bottom), indicating activity-evoked vascular responses. Largest dilations and strongest neuron–vessel correlations occurred in superficial layers (Figure S7 G and H), though significant correlations persisted in penetrating arterioles of deeper layers (e.g., line 9, Figure S7I in supplementary file 1).

### High-speed hemodynamics mapping with tunable spatiotemporal resolutions

The capacity to spatially map rapid blood flow across large brain volumes has broad-reaching implications(Anderle et al., 2025; Iadecola, 2004; Kisler et al., 2017; Sweeney et al., 2018). When applied to hemodynamics imaging, tBessel-TPFM offers two key advantages: robustness against axial motion artifacts and capability to track blood flows in vessels at steep angles to the focal plane. First, a critical challenge in conventional TPFM imaging of awake mice is motion artifacts. While lateral motion artifacts can be readily corrected computationally, axial motion—typically with amplitudes of several microns—can shift structures out of the Gaussian focal plane (Figure S8A). We performed 2D time-lapse imaging 60-μm below pia over a 400×400-μm^2^ FOV at 30-Hz (Figure 3A). A kymograph of a representative image column shows sporadic but high-amplitude fluctuations, characteristic of axial motion artifacts (Figure S8B in supplementary file 1). To further illustrate these artifacts, we color-coded images acquired at different time points across a 20-s period and combined them, revealing substantial structural shifts in and out of the Gaussian focal plane (Figure S8C in supplementary file 1). In contrast, axial motion introduces minimum artifacts in tBessel-TPFM due to the extended axial beam length (Figure S8D in supplementary file 1). We imaged the same FOV with a 0.4 NA Bessel beam at 30-Hz and found little fluctuations from the kymograph (Figure S8E and video S3 in supplementary file 1). Likewise, the color-coded time-lapse images exhibit no noticeable structural shifts (Figure S8F in supplementary file 1).

**Figure 3:**
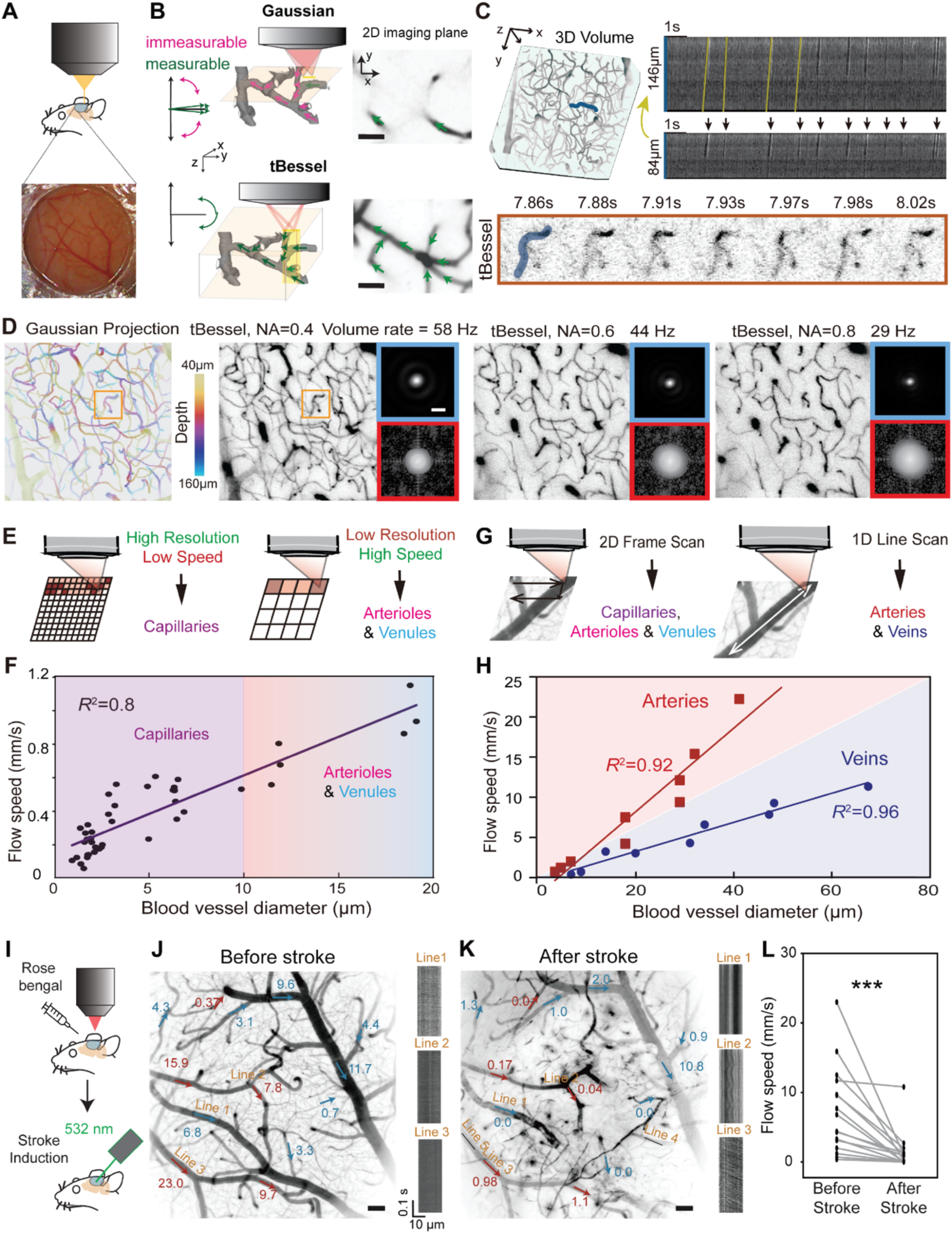
High-speed blood flow mapping in normal and ischemic stroke mice. (**A**) Widefield and schematic view of mice brain vasculature. (**B**) Schematics (left) and images (right) comparing Gaussian (top; 2-μm depth of focus) and tBessel (bottom; 80-μm long) scanning. Gaussian imaging restricts measurements to vessels nearly parallel to the imaging plane, whereas tBessel enables blood flow measurement across a broad range of vessel orientations. Scale bars: 20-μm. (**C**) Workflow for blood flow measurement. Left: 3D Gaussian stack highlighting a vessel segment (blue). Bottom: real-time tBessel imaging showing moving blood cells. Right: kymographs, with blood cells appearing as dark streaks (arrows). Velocities are extracted from 2D tBessel streak slopes and corrected for vessel 3D length. (**D**) Mean intensity projection of a Gaussian stack (400×400×120-μm^3^, 40–160-μm below pia, depth color-coded) versus tBessel scans with NA = 0.4, 0.6, and 0.8. Insets: experimentally measured point spread functions (PSFs, scale bar: 1.5-μm) and optical transfer functions (OTFs). (**E**) Trade-off between spatial resolution and imaging speed: high resolution for capillaries versus low resolution for larger vessels. (**F**) Blood flow speed versus vessel diameter for 42 segments imaged with 0.4 NA tBessel at 58-Hz. (**G**) Frame scanning versus line scanning: restricting scans to vessel regions improves speed and accuracy for large vessels. (**H**) Blood flow speed versus vessel diameter measured with 1 kHz line scans. Red: arteries; blue: veins. (**I**) Schematics of stroke induction. (**J**) tBessel-TPFM hemodynamics of a 1,400×1,400×140-μm³ volume before stroke induction. Blood flow speeds (mm/s) measured by 1 kHz line scans; numbers indicate speed, arrows indicate direction (red: arterioles, blue: veins). Representative kymographs of three linecuts shown at right. Scale bar: 100-μm. (**K**) Same measurement after stroke induction. Scale bar: 100-μm. (**L**) Paired plot of blood flow speed before and after stroke induction (N = 17). Wilcoxon test, p = 1×10^−5^.

Next, we demonstrated that tBessel-TPFM enables tracking of blood flows in vessels that are inaccessible by conventional Gaussian-TPFM. With Gaussian beams, only vessels nearly parallel to the focal plane provide segments long enough within its ∼2-μm depth of focus to measure flow. For instance, a 15-μm segment is required to resolve flows up to 15-mm/s, restricting usable orientations to within ∼7.6° of the focal plane (Figure S8G in supplementary file 1). Analysis of cortical vasculature orientation in mice showed a relatively uniform distribution (Figure S8H in supplementary file 1), meaning Gaussian scanning excludes ∼90% of vessels (Figure 3B and Figure S8I in supplementary file 1). By contrast, our tBessel-TPFM with extended axial beam length can capture vessels over a broad orientation range (Figure 3B). Simulations confirmed that a 60-μm Bessel beam can measure flows up to 1.5 mm/s in vessels angled up to 80° (Figure S8L and Supplementary Note 3 in supplementary file 1).

To assess tBessel-TPFM performance, we imaged Rhodamine B–labeled vasculature in a 400×400×120-μm^3^ cortical volume (Figure 3C). With NA = 0.4 at 58-Hz, unlabeled red blood cells show up as dark streaks in kymographs (Figure 3C, Figure S8J and video S4 in supplementary file 1). To account for the projection nature of Bessel-TPFM, we manually traced the blood vessel segments in 3D Gaussian structural data and measured their 3D lengths (Fan et al., 2020) (Figure 3C, right panels; Methods). Acquisition frame rate sets the upper bound of measurable flow speeds. In tBessel-TPFM, lowering NA enables faster imaging but at a reduced resolution. We quantified this resolution–speed trade-off by imaging the same region with NA = 0.4, 0.6, and 0.8. NA = 0.4 enabled 58-Hz imaging with coarser resolution as shown by the point spread function (PSF) and optical transfer function (OTF), whereas NA = 0.8 provided higher resolution at 29-Hz (Figure 3D). Practically, high-NA, slower scans best capture capillaries, while low-NA, faster scans suit arterioles and venules (Figure 3E). By balancing resolution with acquisition speed using our tBessel module, we mapped 3D flow in 42 vessels (Figure 3F) and confirmed that blood flow speeds are positively correlated with vessel diameters.

### Tracking blood flows before and after ischemic stroke at kilohertz rate

To be able to track individual red blood cells, spatial resolution cannot be reduced indefinitely. Therefore, while tBessel-TPFM offers 10-100× speedup over Gaussian-TPFM in volumetric imaging, the maximum measurable flow speed is still limited by the 2D frame scanning rate, typically tens frame per second (fps) (Figure S8K). To measure ultrafast hemodynamics in large-diameter veins and arterioles, we implemented 1D line scanning at kilohertz rates (Figure 3G)(Meng et al., 2022). In this mode, synchronized galvo–galvo scanning drives the beam back and forth along targeted vessel segments, directly generating kymographs (Figure S9A). To validate this approach, we compared flow speeds measured by 1D line-scan Bessel and Gaussian beams on the same vessel (∼20-μm horizontal span). No significant differences were observed (Figure S9 B and C). Using 1D line scanning tBessel-TPFM, we measured speeds up to 23-mm/s in arterioles and 11.7-mm/s in veins—20× beyond the limits of frame scanning (Figure 3H). Notably, for vessels of similar diameter, flow in arterioles was three times faster than in veins.

Last, we applied 1D line scanning tBessel-TPFM to an ischemic stroke model in mice to investigate hemodynamic changes before and after photothrombosis(Labat-gest and Tomasi, 2013). Rose Bengal was administered intraperitoneally and activated by targeted laser illumination that induced endothelial damage and thrombosis (Figure 3I; Material and Methods). Laser speckle imaging confirmed a sharp reduction of blood flow in the targeted cortical region (∼100-μm diameter), consistent with established ischemic injury (Figure S10 in supplementary file 1). We mapped flow speeds of 17 vessel segments across a 1,400×1,400×140-μm^3^ cortical volume before (Figure 2J) and after stroke induction (Figure 3K). To ensure accurate kymograph construction, 3D Gaussian stacks were acquired before and after each session to extract vessel lengths (Figure S9 E to J in supplementary file 1; Material and Methods). Median flow speeds over 5-s were calculated for each vessel. Before stroke, flows up to 23.0-mm/s were observed. After stroke, nearly all vessels exhibited dramatic reductions, in some cases up to 100-fold (Figure 3L). These results demonstrate that tBessel-TPFM, with tunable resolution and high-speed line scanning, provides a versatile platform for mapping flow dynamics across diverse vessel sizes and orientations in both healthy and pathological states.

### Volumetric neuronal activity imaging with targeted optogenetic stimulation

Neuroscience increasingly relies on experiments that integrate targeted optical perturbations (e.g., optogenetic stimulation or microsurgical ablation) with high-speed volumetric imaging to investigate local and network-level responses. A critical requirement for such paradigm is precise spatial co-registration between the perturbation beam and the imaging volumes (Figure 4A). Existing Bessel modules struggle to meet this requirement because tuning beam parameters shifts the axial center of the imaging Bessel beam relative to the perturbation Gaussian focus (Figure S1 in supplementary file 1), causing misalignment between perturbation and imaging paths. In contrast, our tBessel module resolves this challenge by maintaining a fixed axial center during beam tuning, ensuring that the Bessel imaging volume remains aligned with the Gaussian stimulation beam over a wide range of parameters (Figure 4B). Some mild asymmetry observed in the axial PSF is attributed to the tBessel module, but rather to non-uniformity in the input Gaussian beam (Supplementary note 2 in supplementary file 1). To demonstrate this advantage, we combined two-photon optogenetic stimulation with volumetric calcium imaging. A Gaussian 1040-nm beam delivered spiral-pattern stimulation to a ChRmine-expressing neuron, while a 920-nm tBessel beam simultaneously captured calcium dynamics across a 196×196×80-µm^3^ cortical volume spanning planes above and below the stimulation site (Figure 4C and Figure S11 in supplementary file 1; Material and Methods). A Gaussian reference stack was first acquired to select the stimulation plane and calibrate beam alignment. Optogenetic stimulation reliably evoked robust calcium rises in the targeted neuron (Figure 4D, neuron 1), confirming precise activation. Importantly, calcium spikes were also observed in neurons located above and below the focal plane. (Figure 4D), reflecting functional network responses that would be missed by conventional Gaussian-based imaging. Cross-correlation analysis further revealed putative connectivity among these neurons (Figure 4E), highlighting the ability of tBessel-TPFM to link targeted perturbations with volumetric network-level readouts *in vivo*.

**Figure 4:**
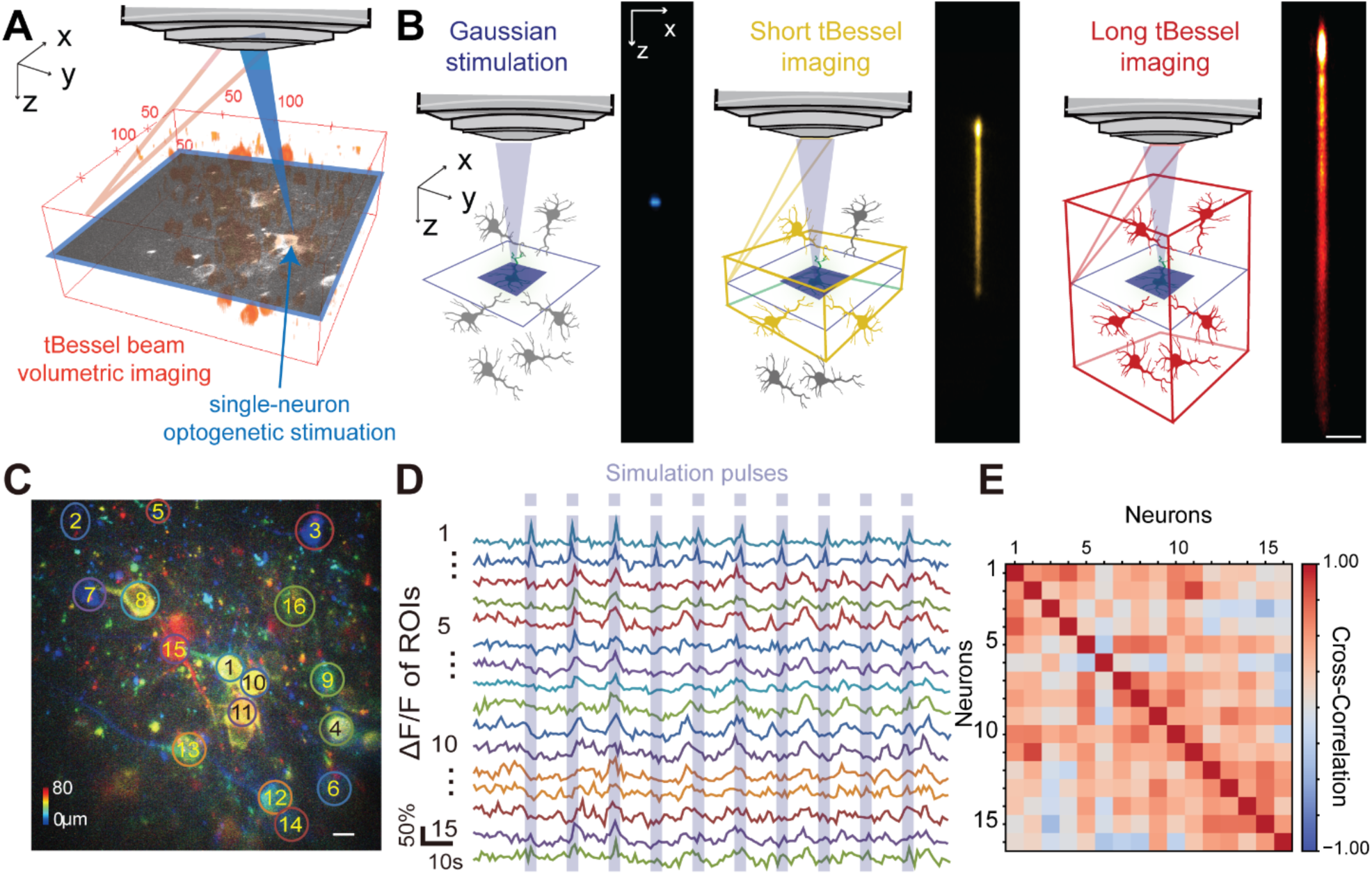
Video-rate volumetric functional imaging with targeted optogenetic stimulation. (**A**) Experimental setup. A Gaussian beam (blue) delivers 3D localized optogenetic stimulation to a neuron, while a tBessel beam enables volumetric imaging symmetrically above and below the stimulation plane. (**B**) Schematics and axial PSFs of the stimulation Gaussian beam (blue), and short (yellow) and long (red) tBessel imaging beams, showing that the extended projection is always symmetric with respect to the Gaussian stimulation plane. The non-uniformity in intensity comes from input Gaussian beam. Scale bar: 10-μm. (**C**) Depth color-coded projection of volumetric functional imaging showing the stimulated neuron (neuron 1) and surrounding neurons; 16 neurons were selected for analysis. Scale bar: 10-μm. (**D**) Representative calcium transients from neurons numbered in (**C**). Light blue bars mark the timing of optogenetic stimulation pulses delivered to neuron 1. (**E**) Pearson correlation map of calcium dynamics among the neurons numbered in (**C**).

### Rapid microglial dynamics imaging with laser ablation

Last, we applied tBessel-TPFM to monitor neuroimmune responses following targeted single-cell ablation. The vasculatures of transgenic mice expressing GFP in microglia was counter-labeled with dextran–Rhodamine B (Figure 5A; Material and Methods). A 3D Gaussian reference stack (200× 200×120-µm^3^) was first acquired, and a single plane was selected for two-photon ablation of a single microglial cell (Figure 5 B and C). Post-ablation, volumetric imaging was performed with a 120-µm-long, 0.7 NA Bessel beam at 15 Hz for 10 min to capture microglial process dynamics (Figure 5D). To maximize interpretability while minimizing photodamage, we imaged the microglia dynamics with low signal-to-noise-ratio (SNR) barely sufficient to identify the rapid extension and retraction of microglial processes dynamics. We then built and trained a neural network based on the recently developed content-aware image restoration (CARE) method to denoise the low SNR images(Weigert et al., 2018) (Figure S9 A to C in supplementary file 1; Material and Methods), followed by Richardson–Lucy deconvolution (Figure S9D in supplementary file 1).

**Figure 5:**
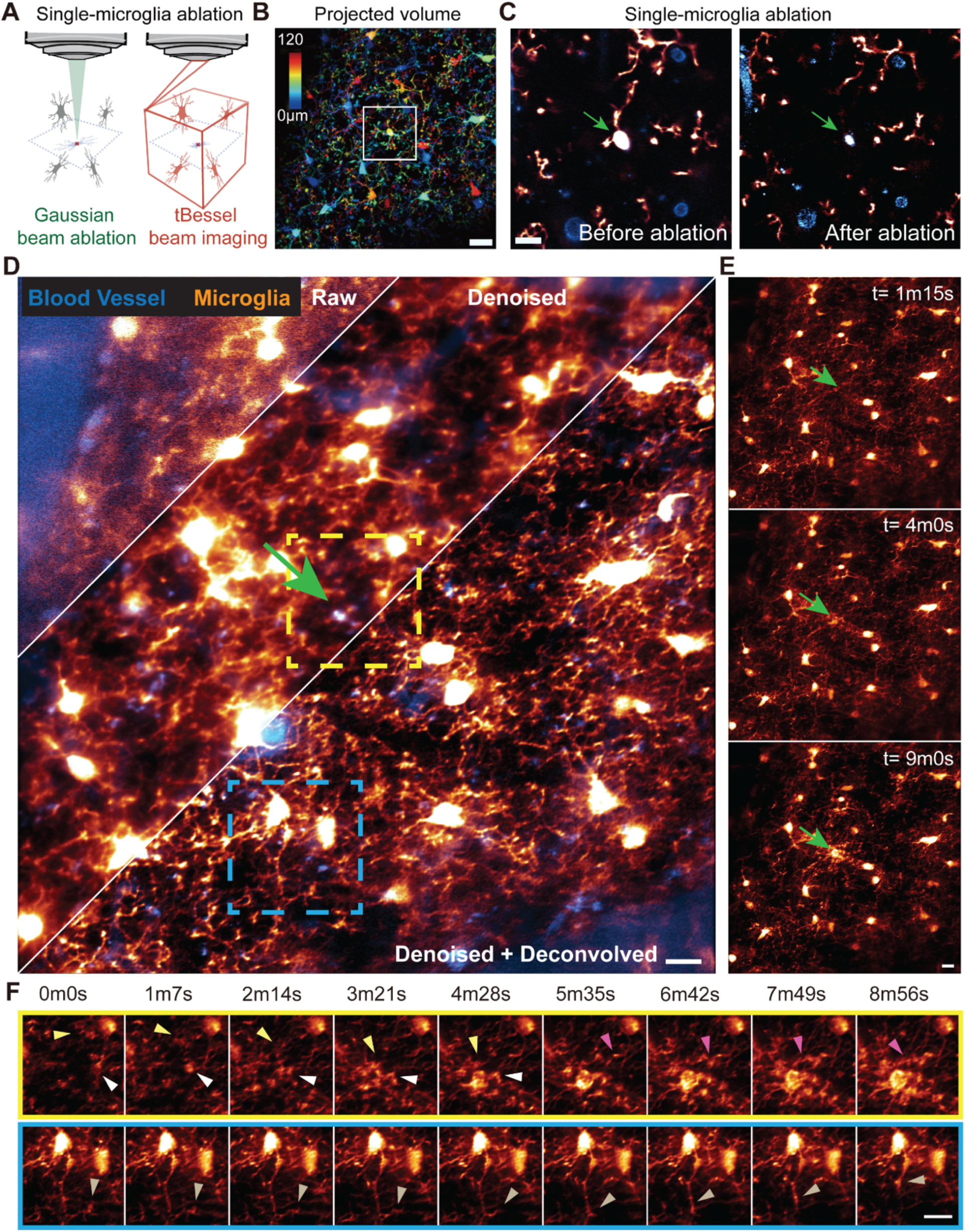
Rapid volumetric imaging of microglial responses to targeted ablation. (**A**) Experimental setup: a Gaussian beam (green) delivers targeted ablation of a single microglial cell, while a tBessel beam (red) provides symmetric volumetric imaging above and below the ablation plane. (**B**) Mean intensity projection of a 200×200×120-μm^3^ Gaussian stack, depth color-coded. White box marks the targeted microglia. Scale bar: 20-μm. (**C**) Gaussian plane images of the targeted cell before (left, arrow) and after (right, arrow) ablation. Scale bar: 10-μm. (**D**) Representative 15 Hz volumetric tBessel data with a 120-μm-long beam (NA = 0.7). Raw (left), denoised (center), and deconvolved (right) images. Scale bar: 10-μm. (**E**) Time-lapse snapshots showing process extension toward the ablation site (arrowheads) at 1m15s, 4m0s, and 9m0s, revealing a two-wave response. Scale bar: 10-μm. (**F**) Zoomed views of two regions (green and blue boxes in (**D**)). Arrowheads indicate process extension toward the ablation site (top, green box) and retraction opposite the lesion (bottom, blue box) across the 10-min imaging period. Scale bar: 5-μm.

Following ablation, neighboring microglia exhibited rapid responses: their somata remained stationary, but processes extended dynamically toward the damage site. Notably, two distinct waves of process extension were observed, the first arriving around 4 minutes post-ablation and the second at approximately 9 minutes (Figure 5E and video S5 in supplementary file 1). Detailed analysis of process dynamics revealed that microglia not only extended processes toward the injury (Figure 5F, top row) but also retracted processes opposite to the damage site (Figure 5F, bottom row). Furthermore, the advancing processes displayed exploratory movements, probing the surrounding environment rather than following linear trajectories. While prior work has shown that microglia extend processes toward injury sites without soma migration(Davalos et al., 2005), these fine-scale dynamics—including two-wave responses, coordinated extension-retraction patterns, and rapid local probing—have not been previously reported, as prior technologies lacked sufficient temporal resolution for such observations.

## Discussion

In summary, we developed tBessel-TPFM, a compact and versatile platform for high-speed volumetric imaging of brain dynamics with independently adjustable spatial resolution, axial projection length, and stable focal positioning. The tBessel module enables adaptive tuning of the Bessel beam’s resolution, imaging depth, and side-ring excitation, optimizing performance for the heterogeneous vascular structures and dynamic processes found across brain regions and depths. Critically, it addresses two major barriers to the broader adoption of Bessel beam TPFM: the complexity of existing implementations and the lack of tunability. In addition to providing full control over the Bessel beam profile, the module features a light-efficient design—achieved without diffractive elements or photomasks—and requires minimal instrumentation and programming, making it accessible to a wide range of users.

Unlike conventional systems that rely on costly programmable SLMs(Lu et al., 2017) or passive optics that either lack tunability or introduce undesired axial focus shifts of the Bessel beam(Dickey and Conner, 2011; Lu et al., 2018; Takanezawa et al., 2021), tBessel-TPFM achieves full optical tunability without focus drift using only off-the-shelf refractive components. This adaptability allows researchers to match imaging parameters to specific structures—ranging from fine microglial processes and capillaries to large vessels and neuronal ensembles—within a single platform. The center-stable beam design further enables integration with optical perturbation techniques, including optogenetic stimulation and laser ablation, where precise co-registration between imaging and stimulation beams is essential.

We demonstrated the broad applicability of tBessel-TPFM for brain imaging across several challenging applications. Importantly, the center-stable tunability of our tBessel-TPFM enables biological observations that are difficult or impossible to obtain with conventional Gaussian-TPFM or prior Bessel-TPFM implementations. First, by optimizing the Bessel foci length in accordance with the brain vascular structures and distribution, we achieved volumetric imaging across millimeter-scale cortical volumes at sub-capillary resolution. These observations enable quantitative measurements of blood flow speeds in microcapillaries, arterioles and venules as well as direct observations of NVC across layers in awake mice. The extended Bessel projection, combined with focal jumping and lateral tiling, allowed rapid mapping of blood flow and vessel diameter changes over cortical volume. In addition, high-speed Bessel line scans at kilohertz rates captured ultrafast blood flow in large, steeply oriented vessels before and after ischemic stroke induction. Moreover, the platform supported precise targeted optogenetics with volumetric readout for volumetric neural connectivity mapping. Because the axial center of the Bessel beam remains fixed during NA adjustments, we achieved stable co-alignment of volumetric imaging and holographic optogenetic stimulation for all-optical physiology, enabling reliable multi-plane readout of light-evoked calcium transients. Furthermore, we performed rapid tracking of microglial process dynamics after single-cell-level laser ablation, revealing fine-scale two-phase response patterns across ∼100 µm depths that would be difficult to capture with low-speed Gaussian-TPFM.

Looking ahead, future developments could include closed-loop automation, in which real-time image analysis guides on-the-fly optimization of Bessel beam parameters. In addition, integration with three-photon excitation(Wang and Xu, 2020) and adaptive optics(Hampson et al., 2021) would extend high-resolution brain imaging into subcortical regions. Furthermore, advances in optical engineering—such as novel nondiffractive beam designs—may further mitigate the trade-offs among spatial resolution, axial coverage, and side-ring excitation inherent to Bessel beams(Chen et al., 2024). Nevertheless, even in its current form, the demonstrated capabilities of tBessel-TPFM already open substantial opportunities for both basic and translational brain science, including studies of vascular and immune function in neurodegenerative disease models, investigations of neurovascular and neuroimmune coupling, and the development of targeted therapeutic strategies. Beyond two-photon microscopy, the tBessel module’s versatile tunability makes it readily adaptable to other Bessel-based modalities, including light sheet microscopy(Planchon et al., 2011), optical tweezers(Garcés-Chávez et al., 2002), and ophthalmology(Khonina et al., 2020).

## Materials and Methods

### Animal use

All experimental protocols followed the National Institutes of Health guidelines for animal research. All procedures involving mice were approved by the Institutional Animal Care and Use Committee at the University of Maryland School of Medicine. Male and female mice (C57BL/6J, stock #000664; CX3CR1-GFP, stock #005582; Jackson Laboratories), aged two months or older, were used in this study. Mice were housed in groups of 1–5 per cage under a normal light cycle prior to surgery.

### Assembly and incorporation of the tBessel module in TPFM

The tBessel module consists of three axicons, three lenses, and one iris to enable independent control of the numerical aperture (NA) and the energy confinement of the generated Bessel beam. The first axicon (AX252B, Thorlabs) converts the incoming collimated illumination beam into a Bessel beam. An iris (SM2D25D, Thorlabs) is positioned at the minimum beam waist—corresponding to the sample-conjugated plane—to tune the beam’s energy confinement. A plano-convex lens (AC254-125-B, Thorlabs) is placed such that its front focal plane aligns with the iris. Next, a matched pair of axicons with identical angles (AX255B, Thorlabs) is used to adjust the NA of the Bessel beam. Finally, a 4f lens pair (AC254-125-B and AC254-150-B, Thorlabs) relays the virtual ring focus onto a galvo scanner conjugated to the objective’s back focal plane. The entire tBessel module can be integrated into any commercial TPFM via two mirrors mounted on a rail assembly, allowing rapid switching between Gaussian and Bessel beam modes. Imaging was performed on a custom-built two-photon microscope (Modular In vivo Multiphoton Microscopy System, MIMMS) designed by Janelia Research Campus, Howard Hughes Medical Institute. In Gaussian mode, after beam expansion, the excitation laser passes through a Pockels cell and beam expander. The tBessel module was installed as a switchable unit using a linear actuator between the resonant scanner and a preceding mirror in the conventional laser path. In both modes, the beam is focused onto a resonant scanner (8 kHz line rate) for x-axis scanning, followed by a Y-galvo mirror for y-axis scanning. The Y-galvo is optically conjugated by a pupil relay to the objective’s back focal plane.

### Stereotaxic surgery for *in vivo* imaging

Surgeries were performed using a stereotaxic apparatus (RWD) and aseptic technique as previously described(Lu et al., 2020, 2017; Zhang et al., 2023). Mice were anaesthetized with isoflurane (1–2% by volume in O_2_). A craniotomy was made over the left V1 with the dura left intact. For mice that required virus injection, a beveled glass micropipette (15–20 μm tip, Drummond Scientific) was back-filled with mineral oil and connected to a hydraulic manipulator (Narishige, MO10) for precise delivery. AAV viral solutions (AAV2/1.syn-GCaMP7s, 9 × 10^12^ GC/mL; Plasmid #104487 or AAV8-hSyn-GCaMP6m-p2A-ChRmine-Kv2.1, 2 × 10¹² vg/mL; Plasmid #131004, Addgene) were injected at depths of ∼200–400 μm below the pia at coordinates Bregma −3.4-mm and −4.0-mm (AP) and 2.2-mm and 2.6-mm (ML), delivering 10–20-nL over 2-minutes per site. For all animals, following injection or craniotomy, a no. 1.5 glass coverslip (Fisher Scientific) was embedded in the opening and sealed with dental acrylic. A titanium headpost was attached with cyanoacrylate glue and dental acrylic. Mice recovered for 3–4 weeks to allow opsin and GCaMP expression.

### Blood vessel labeling with fluorescent dyes

For vascular imaging, 50-μl of 2% dextran conjugated Rhodamine B (70-kDa) or 5% dextran-conjugated fluorescein (70-kDa) was injected retro-orbitally in mice anesthetized with isoflurane (maintained at 1–2% by volume in air).

### Stroke induction

Focal ischemic stroke was induced via photothrombosis using Rose Bengal. A cranial window over V1 was implanted at least three weeks before stroke induction. Mice were anesthetized with 1.5% isoflurane and kept on a 37 °C heating pad. Rose Bengal solution (50-μL, 20-mg/mL in PBS) was injected retro-orbitally. One minute later, a green laser (532 nm, 3-mW) illuminated a cortical region away from major vessels for 5 minutes. Ischemic core and peri-infarct areas were identified by subsequent imaging.

### *In Vivo* two-photon imaging

Gaussian or Bessel beams were scanned using a galvo-resonant mirror for 2D imaging and galvo–galvo scanners for line scans. Fluorophores were excited using a femtosecond laser (Chameleon Discovery, Coherent) and imaged with either a 25×, 1.05 NA water-immersion objective or a 16×, 0.8 NA Nikon water-immersion objective (Olympus). Emitted photons were reflected by a dichroic mirror (FF705-Di01, Semrock) and then separated by a dichroic mirror (565DCXR, Chroma). Green and red channels were detected by PMTs (H16201P-40//004, Hamamatsu) after filtering with a 510/84 nm bandpass (green; Edmund 84-097) and two 750SP short-pass filters (red; Edmund 64-332). Images were acquired with ScanImage(Pologruto et al., 2003).

### Line-scan analysis of blood flow speed

Blood flow speed was determined from red blood cells displacement between sequential line scans using cross-correlation, as described previously(Kim et al., 2012). The pixel shift was converted to microns, and velocity calculated as *v* =Δ*x/*Δ*t*, where Δ*t* is the time between line scans. For vessels not perpendicular to the imaging plane, a five-step geometric correction was applied: 1. Overlay the Bessel beam scan location onto Gaussian point scan stacks to align sessions. 2. Measure the lateral displacement *dx* of the vessel segment. 3. Determine the vertical displacement *dy* by comparing z-slices at each segment endpoint. 4. Calculate vessel tilt angle 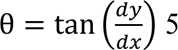. Correct velocity using 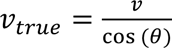. This correction approximates the true flow speed; multiple measurements across the vessel improve accuracy.

### Segmentation of blood vessels and kymographs

Image data were converted to double precision, logarithmically transformed twice (log_10_) to compress dynamic range, smoothed with a Gaussian filter (σ = 1), and thresholded at 0.32–0.36 to produce a binary mask. The binary image was converted to double precision, and pixel counts per row were computed to quantify vessel width over time.

### Visual Stimulation in awake mice

For head-fixed awake imaging, mice were habituated 1 week after surgery. Each session involved restraining the body under a half-cylindrical cover to reduce movement, repeated 3–4 times per mouse for 15–60 min each. Imaging began 3–4 weeks after virus injection. Visual stimuli were presented on a blue-light tablet monitor (SCEPTRE) positioned 15 cm from the right eye (70° × 70° visual field, oriented ∼40° relative to the body axis). Stimuli were generated in MATLAB using Psychophysics Toolbox. Each trial included 8-s of a blank gray screen followed by 8-s of drifting gratings (0.05 cycles/degree, 1-Hz temporal frequency) in eight randomized directions, with five repeats per direction. Imaging (via ScanImage) started 4 s into the blank period, capturing 4 s of gray screen and 4 s of stimulus.

### Simultaneous two-photon optogenetics and calcium imaging

Wild-type C57BL/6J mice (8–12 weeks) previously injected with AAVs for calcium indicator and opsin expression were anesthetized (1–2% isoflurane) and head-fixed under the microscope. Layer 2/3 neurons with clear somata were identified under Gaussian-mode imaging at 920-nm (<50-mW at the sample). A 3D z-stack (∼150-μm, 2-μm step size) was acquired around target cells. Optogenetic stimulation of ChRmine was delivered via the same objective using a 1040 nm spiral scan (logspiral, 5 revolutions, 5 ms duration) to the soma, following established protocols(Daie et al., 2021; Sridharan et al., 2022). 10 stimulations (5-s inter-stimulus interval) were applied at 70–100-mW post-objective. Volumetric imaging was then performed in Bessel mode using the same stimulation protocol.

### Two-photon laser ablation of single microglia

CX3CR1-GFP mice with cranial windows were anesthetized (1–2% isoflurane) and head-fixed. Targeted microglia were identified by soma morphology. A 900-nm laser (∼60-mW) was focused 70-μm below the dura onto the soma center and delivered for 5-s in point-stimulation mode. Successful ablation was confirmed by loss of GFP signal and morphological changes.

### Denoise and deconvolution

To improve the SNR of microglia imaging data under low excitation power, we trained a Content-Aware Image Restoration (CARE) neural network on paired low-SNR and high-SNR images. Training data were generated from time-lapse image sequences of microglia acquired at various cortical locations under identical imaging conditions. For each location, low-SNR inputs were single frames extracted from the video, while high-SNR ground truth images were generated by averaging ten consecutive frames from the same sequence. These paired datasets were used to train the CARE model with a U-Net architecture. After training, the network was applied to denoise time-lapse datasets acquired during microglial responses to targeted ablation. Denoised images were subsequently processed by 20 iterations of Richardson–Lucy deconvolution.

### Statistical analysis

Analysis was performed using MATLAB with standard functions and custom scripts. Data normality was assessed; parametric tests were applied to normally distributed data, and nonparametric tests to others. For normal data, results are reported as mean ± SEM with bar graphs; for non-normal data, as median ± IQR with box plots (Tukey whiskers ±1.5×IQR). Paired nonparametric data were tested with the Wilcoxon signed-rank test; unpaired nonparametric comparisons used the Wilcoxon rank-sum test. Statistical significance thresholds were: *P < 0.05, **P < 0.01, ***P < 0.001. Experiments were not blinded. Sample sizes were chosen based on common practice in the field.

## Acknowledgments

We thank F. Liu, X. Zhang, and J. Wu for helpful discussions.

## Funding

National Institutes of Health 1R21EB035681 (MJL, YQ, TMF)

Princeton Alliance for Collaborative Research and Innovation (TMF)

Princeton School of Engineering and Applied Science Innovation Grants (MJL, TMF)

National Institutes of Health R21AG077631, R03NS123733, R03NS128459, and R21AG074978 (JW, MW, CR, YL)

Maryland Stem Cell Research Fund 2024-MSCRFD-6363 and 2022-MSCRFL-5893 (JW, MW, CR, YL)

Maryland Stem Cell Research Fund 2022-MSCRFD-5886 (PW)

National Institutes of Health R01DA056739 (PW)

Maryland Stem Cell Research Fund Postdoc Fellowship 2024-MSCRFF-6328 (JW)

## Author contributions

Conceptualization and supervision: T.-M.F. and Y.L.

Microscope design, built, and alignment: M.J.L. and T.-M.F.

Theory and simulation: M.J.L. and Y.Q.

Mouse surgery, stroke, and imaging: Y.L., J.W. M.L., and P.W.

Data analysis and visualization: M.J.L., M.W., C. R.

Funding acquisition: T.-M.F., Y.L., P.W., and J.W.

Figures and videos: M.J.L., Y.L., and T.-M.F.

Writing: M.J.L., Y.L., and T.-M.F. with input from all authors.

## Competing interests

MJ.L., Y.L., and T.-M.F. are on a patent filed by Princeton University related to this work.

## Data and materials availability

Interested individual can request all datasets directly from the corresponding authors.

