## Supplementary Materials for "Tunable Bessel beam two-photon fluorescence microscopy for high-speed volumetric imaging of brain dynamics"

### Supplementary Text

#### Supplementary Note 1: Theoretical analysis of tunable Bessel module

The tunable Bessel (tBessel) module generates a Bessel beam with independently tunable numerical aperture (NA) and lateral confinement (ΔNA), while maintaining a fixed axial beam center. This is achieved through a modular optical design composed of standard refractive components.

A collimated Gaussian beam first passes through an axicon-lens pair (A1 and L1), where the first axicon (A1) imparts a conical phase to the beam, generating diverging rays at a fixed angle $\theta = (\mathrm{n} - 1)\alpha$ that form the Bessel beam, whose shape is determined by the axicon's apex angle $\alpha$ and refractive index n. These rays are then focused by lens L1 into a ring-shaped intensity profile (Bessel ring focus) at a pupil-conjugate plane.

This focused ring is then relayed through a symmetric pair of identical axicons (A2 and A3), which together form an angle-preserving optical system. In such a configuration, rays entering at a given angle exit at the same angle independent of the axicon-pair separation. This angle-preserving property is central to the tunability of our Bessel module. While the propagation angles of the rays remain fixed, changing the separation between the axicons shifts the radial position at which the rays exit, thereby linearly altering the radius of the ring formed at the pupil plane. As a result, increasing the axicon separation effectively increases the NA of the Bessel beam. Crucially, the symmetric geometry of the axicon pair ensures that this adjustment introduces minimal phase curvature, maintaining the axial position of the beam's center. The exiting light rays can be traced back to a virtual ring focus, which is then imaged onto the scanning galvo-mirrors (pupil conjugate) via a 4f relay formed by lenses L2 and L3.

To control ΔNA, and therefore the lateral confinement of the Bessel beam, an iris with adjustable opening size is introduced at a sample-conjugate plane. A smaller iris opening increases the ΔNA, leading to a shorter Bessel beam with tighter energy confinement and reduced side-lobe excitation. Conversely, a larger iris opening reduces the ΔNA, resulting in a longer Bessel beam with more pronounced side-lobe excitation. The iris can theoretically be placed at any sample-conjugate planes. In our implementation, the iris is positioned near the formation of the initial Bessel beam, immediately after A1, offering stable and efficient control over beam length.

To quantify the effect of the system in tuning NA and ΔNA, we define the input beam diameter as $D$, axicon 1(A1) having angle of $\alpha$, the iris opening diameter as $l$, the lens 1 (L1) have a focal length of $f_1$, the pair of axicon A2-A3 having equal angle of $\beta$, and are separated by $L$, the lens 2 (L2) have a focal length of $f_2$, as shown in Figure S2B. All the axicons have the same refraction index $n$. Geometrical optics analysis gives the following relationships:

1. Control of ΔNA and Bessel beam length (B) via Iris:

$$\Delta NA = 1.22\frac{\lambda f_1}{l} \cdot 2 = 2.44\frac{\lambda f_1}{l}$$

which is the spot size generated by the Bessel beam after the iris clip of opening $l$. The resulting Bessel beam is thus

$$B = \frac{2 \cdot 2.44\lambda}{NA \cdot \Delta NA} f_2^2 = \frac{2l}{NA} \cdot \frac{f_2^2}{f_1}$$

2. Control of NA via axicon pair (A2-A3) separation $L$:

$$NA = 2(L - L_c) \cdot \tan\big(\beta(n-1)\big)$$

where

$$L_c = f_1 \cdot \frac{\tan\big(\alpha(n-1)\big)}{\tan\big(\beta(n-1)\big)} \simeq f_1 \cdot \frac{\alpha}{\beta}$$

The range of $L$ needed is

$$L_c \leq L \leq \frac{NA_{max}}{2 \cdot \tan\big(\beta(n-1)\big)} + L_c$$

Some notes about the system:

1. While the axicon pair A2-A3 can be placed anywhere after L1, our test shows that it is best to place A2 before the focal plane of L1, and place A3 after the focal plane of L1.
2. The distance between the iris and A1 is: $d \simeq \frac{D}{4(n-1)\alpha}$
3. There is an implicit restriction due to the aperture size of the optical component. Assume each component is circular in shape, with diameter $a$. Then, we have the following constraint:

$$2f_1 \tan \alpha(n-1) + l \leq a$$

Notice that the left side is equivalent to the ring from A1 directly to L1, and the largest $l$ that can take is $D/2$, thus the above relation translate to:

$$2(f_1 + d) \tan \alpha(n-1) \leq a$$

This gives us an upper bond of the input beam diameter:

$$D \leq 2a - 4f_1\alpha(n-1)$$

**Supplementary Note 2: Optical alignment**

While the core principles of the tBessel module rely on geometric ray propagation and symmetric beam shaping, several practical considerations in the optical configuration can affect performance. Below, we outline key design choices and their justifications, specifically the orientation choices of axicons, and the optional possibility of a photomask.

The orientation of the first axicon (A1), which imparts a conical phase to the collimated Gaussian input beam, is not critical to the core functionality of the system. In our implementation, we oriented A1 with the conical surface facing the incoming beam. Orienting the flat surface toward the incoming beam is also viable and may help reduce axicon imperfections. Both orientations should yield comparable beam profiles, and the choice can be made based on practical considerations without significantly affecting system performance.

The orientation of the second and third axicons (A2 and A3), which constitute the angle-preserving axicon pair, plays a crucial role in maintaining wavefront flatness and minimizing optical aberrations. These axicons should be arranged such that A2 has its conical surface facing the incoming beam and A3 has its flat surface facing the incoming beam, resulting in the flat surfaces of A2 and A3 facing each other. While either the conical or flat surfaces of the axicon pair could theoretically be arranged to face inward in a symmetric geometry, we have experimentally found that the flat-inward configuration best preserves the phase uniformity. This preference arises from the fact that the beam entering the axicon pair already forms a focus, meaning that peripheral rays remain peripheral after transmission. If the conical surfaces are inward facing, these outer rays traverse a longer optical path between the axicons, leading to accumulated phase shifts and spherical aberrations. Orienting the flat surfaces inward mitigates these effects by equalizing optical path lengths across the beam profile.

Additionally, our tBessel system does not require a corrective photomask, as the Bessel focus generated maintains a flat wavefront across different tuning conditions. This ensures that the Bessel beam remains center-stable without axial drift, in contrast to traditional single-axicon or axicon-lens implementations where wavefront tilt or curvature leads to axial displacement and necessitates photomask correction. Some mild asymmetry observed in the axial PSF across all NA (Figure 4B, Figure S2I), is attributed not to the tBessel module, but rather to non-uniformity in the input Gaussian beam. Specifically, a centrally peaked intensity distribution results in preferential energy deposition on one side of the Bessel beam, which manifests as the axial PSF non-uniformity. Nonetheless, for users who wish to implement additional masking, a corrective photomask can be incorporated by inserting a 4f relay immediately after the A1-L1 pair, with the mask placed at the intermediate Fourier plane. While unnecessary in our system, this optional configuration provides flexibility for highly specialized applications or non-ideal optical setups.

**Supplementary Note 3: Theoretical calculation of maximum measurable blood flow speed**

Consider a blood vessel oriented at an angle $\theta$ relative to the horizontal plane as shown in FigureS8J. Let $B$ denote the length of the Bessel-illuminated region, and $c$ represent the characteristic size of the blood cells under observation. If $t$ is the time interval between the entry and exit of a blood cell through the Bessel-illuminated area, then the maximum measurable blood flow speed is:

$$V_{max} = \frac{l}{t} = \frac{B + c}{\sin\theta} \cdot \frac{1}{t}$$

Normally we would need at least 4 frames to track the blood cell, thus this maximum is further decreased by 4 times.

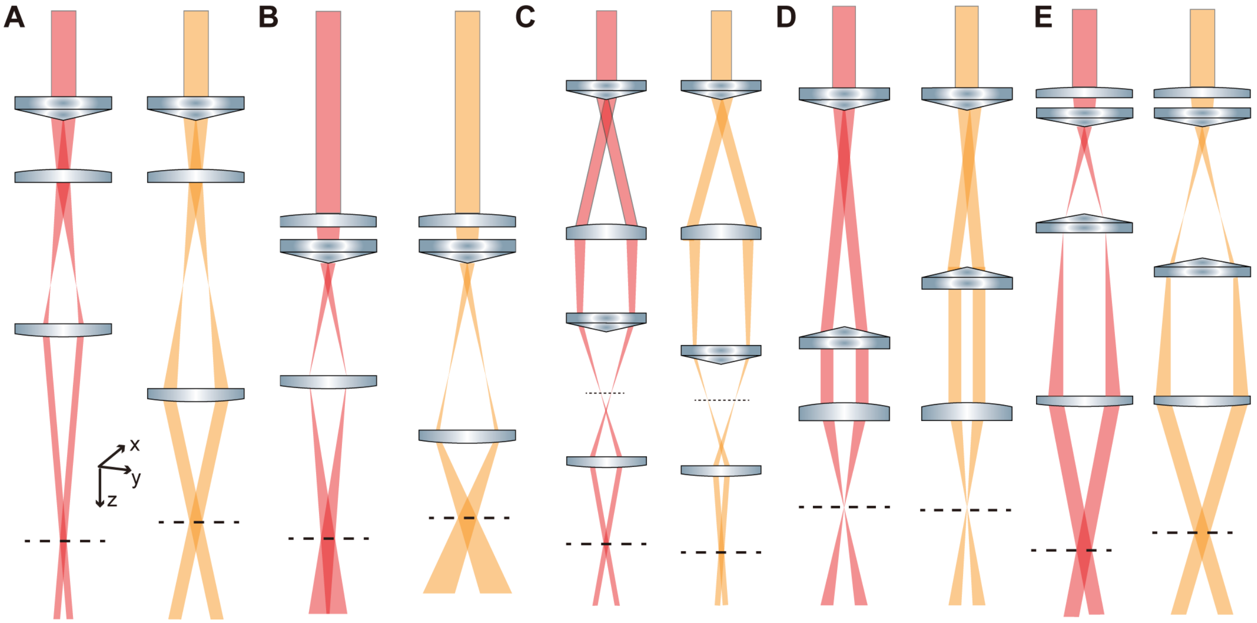


**Figure S1 Comparison of Bessel beams tuning method.** (**A**) Axicon-lens design. This method adjusts the NA and ΔNA through defocusing the last lens. It lacks independent control over NA and ΔNA, and the waist of the generated Bessel beam shift axially when being tuned. (**B**) Lens-axicon design. Similar to (**A**), this method relies on the last lens for tuning NA and ΔNA. It also leads to the generated Bessel beam waist shift axially when being tuned. (**C**) Dual-axicon design. This method provides independent control of NA and ΔNA, but the generated pupil ring has a non-flat wavefront, causing the Bessel beam waist to shift axially. (**D**) Focused Bessel beam generation method utilizing an axicon pair. This approach achieves independent NA and ΔNA control without shifting the Bessel beam waist. However, this method is limited to generate short Bessel beams. (**E**) This design enables independent NA and ΔNA control, but the waist of the generated Bessel beam shift axially when being tuned.

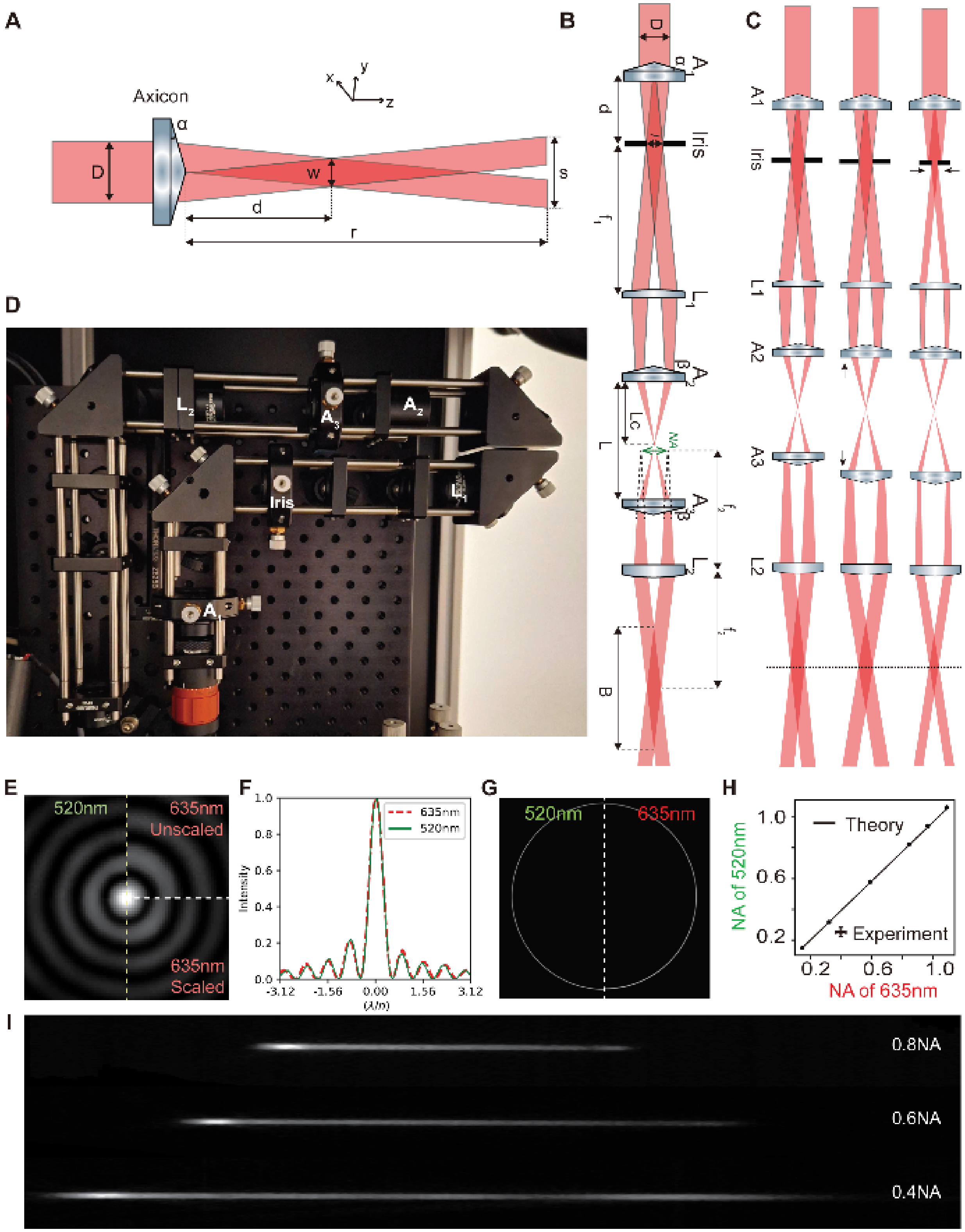


**Figure S2 Tunable Bessel module.** (**A**) Input and output of an axicon. (**B**) Optical diagram of tunable Bessel module with system parameters defined. (**C**) Optical diagrams showing the module’s capability to independently tune the NA (comparing top and middle row) and ΔNA (comparing bottom and middle row). (**D**) Photo of the tunable Bessel module integrated into a Gaussian-TPFM. (**E**) Experimental comparison of simultaneously two-color Bessel beams with 520 nm (left) and 635 nm (top right) laser. Bottom right corner shows the rescaled 635 nm Bessel by wavelength. (**F**) Intensity profiles from the vertical dashed white line of (**E**) comparing the two colors. The horizontal axis is rescheduled by their corresponding wavelengths. (**G**) Experimental comparison of the two colors pupil profile. (**H**) Experimental NA comparison between the two colors at various NAs. (**I**) Experimental axial PSF of NA at 0.4, 0.6, and 0.8 NA. The slight non-uniformities in axial intensity are consistent across different NA settings. This is not induced by the tBessel module itself but rather the centrally peaked intensity distribution of the input Gaussian beam.

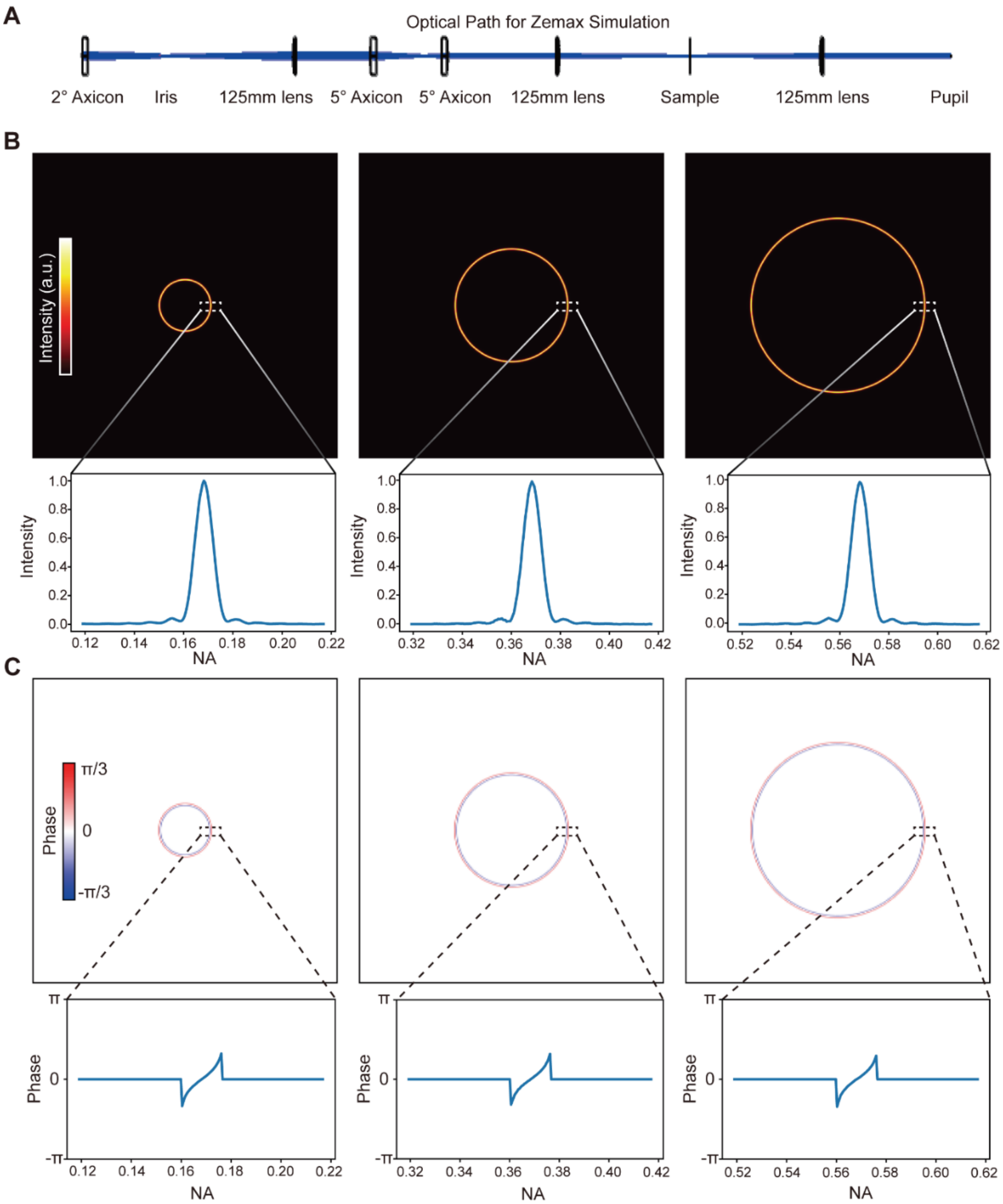


**Figure S3 Zemax simulation of the tunable Bessel module: NA tuning.** (**A**) Zemax diagram of the tunable Bessel module. (**B**) Pupil intensity profiles (top) of three Bessel beams with NA=0.17, 0.37, and 0.57 (same ΔNA) by varying the separation between the two 5° axicons. Line cuts of the intensity profiles (bottom) show that ΔNA remains constant, demonstrating that adjustments to the axicon pair distance do not impact the delta NA of the system. (**C**) Pupil phase profiles (top) and line cuts (bottom) of the three Bessel beams in (**B**).

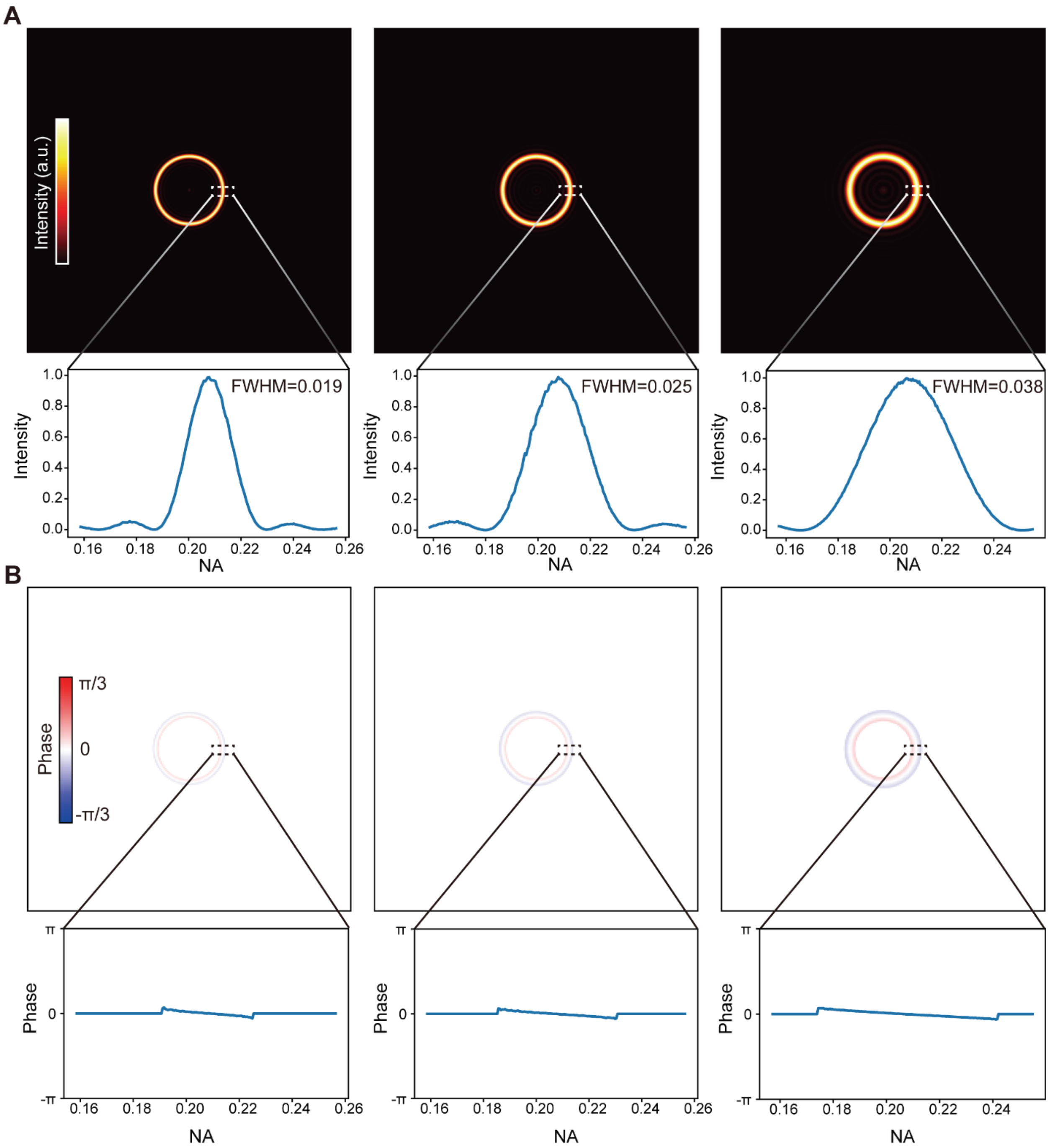


**Figure S4 Zemax simulation of the tunable Bessel module: ΔNA tuning, pupil plane characterization.** (**A**) Pupil intensity profiles (top) and line cuts (bottom) of three Bessel beams with ΔNA=0.019, 0.025, and 0.038 (same NA) by varying the iris opening diameter. (**B**) Pupil phase profiles (top) and line cuts (bottom) of the three Bessel beams in (**A**).

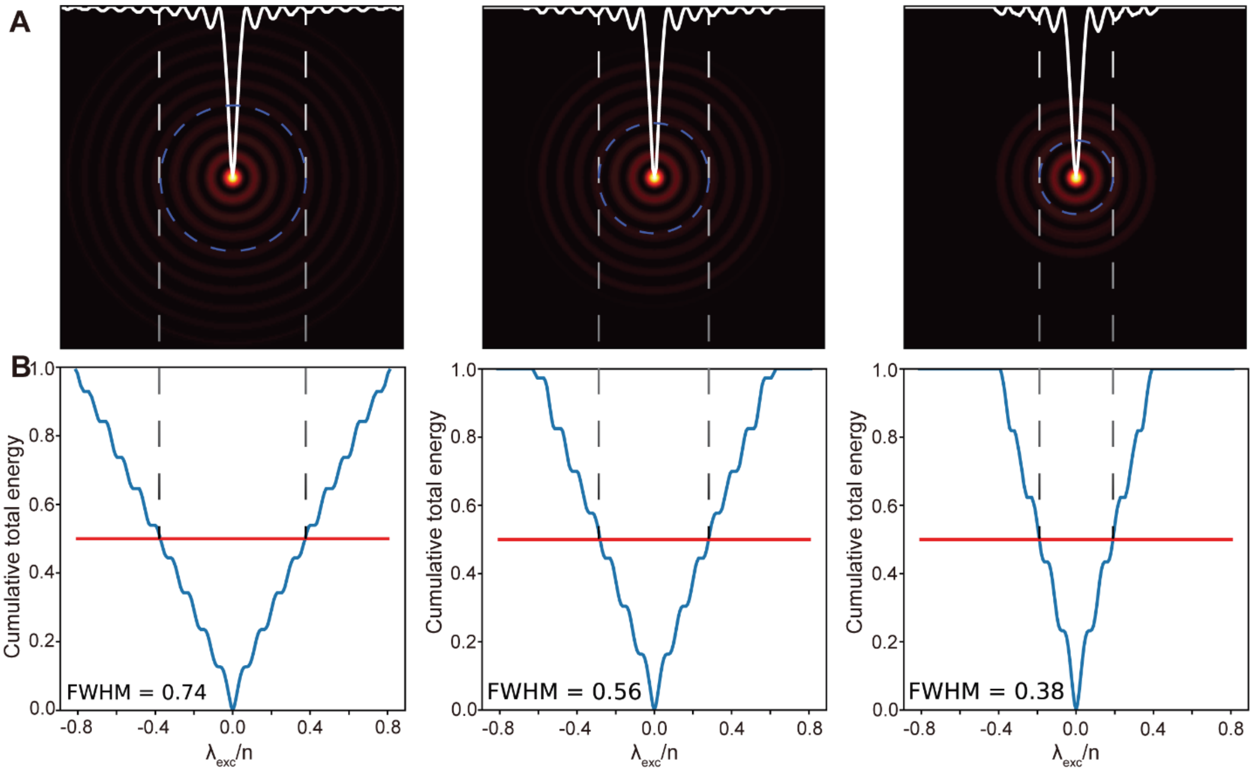


**Figure S5 Zemax simulation of the tunable Bessel module: ΔNA tuning, sample plane.** (**A**) Sample intensity profiles (top) of three Bessel beams with ΔNA=0.019, 0.025, and 0.038 (same NA) by varying the iris opening diameter. The larger the ΔNA, the less side-ring excitation the Bessel beam has. (**B**) Cumulative total energy distribution. The red lines indicate where the half total energy reside.

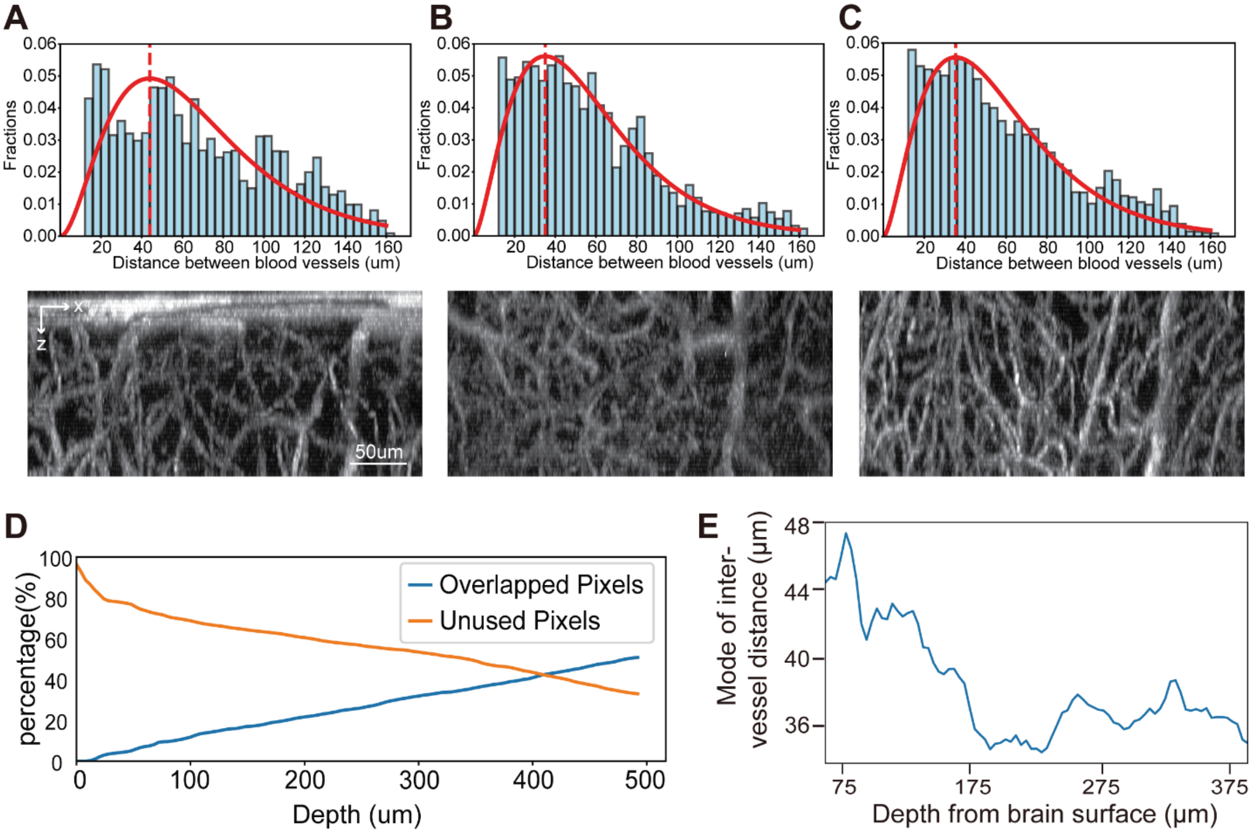


**Figure S6 Analysis of inter-vessel distance at different depths in mouse cortex.** (**A**) Histogram (top) and representative image (bottom) of inter-vessel distance 0–165-µm below pia. The data is fitted to a gamma function, with the mode indicating an optimal Bessel beam length of 44-µm for imaging (red dashed line). (**B**) Histogram (top) and representative image (bottom) 165–330-µm below pia, with a mode of 36-µm (red dashed line), showing slightly closer vessel spacing compared to the surface. (**C**) Histogram (top) and representative image (bottom) 330–500-µm below pia, where the mode of vessel separation is 38-µm (red dashed line). (**D**) Plots illustrating vessel occupancy and overlap along the axial direction from surface to depth. The orange line represents the fraction of pixels in the mean intensity projection that are occupied by blood vessels, while the blue line shows the fraction of these vessel-containing pixels that exhibit overlap. (**E**) Average blood vessel diameter varies at different depths.

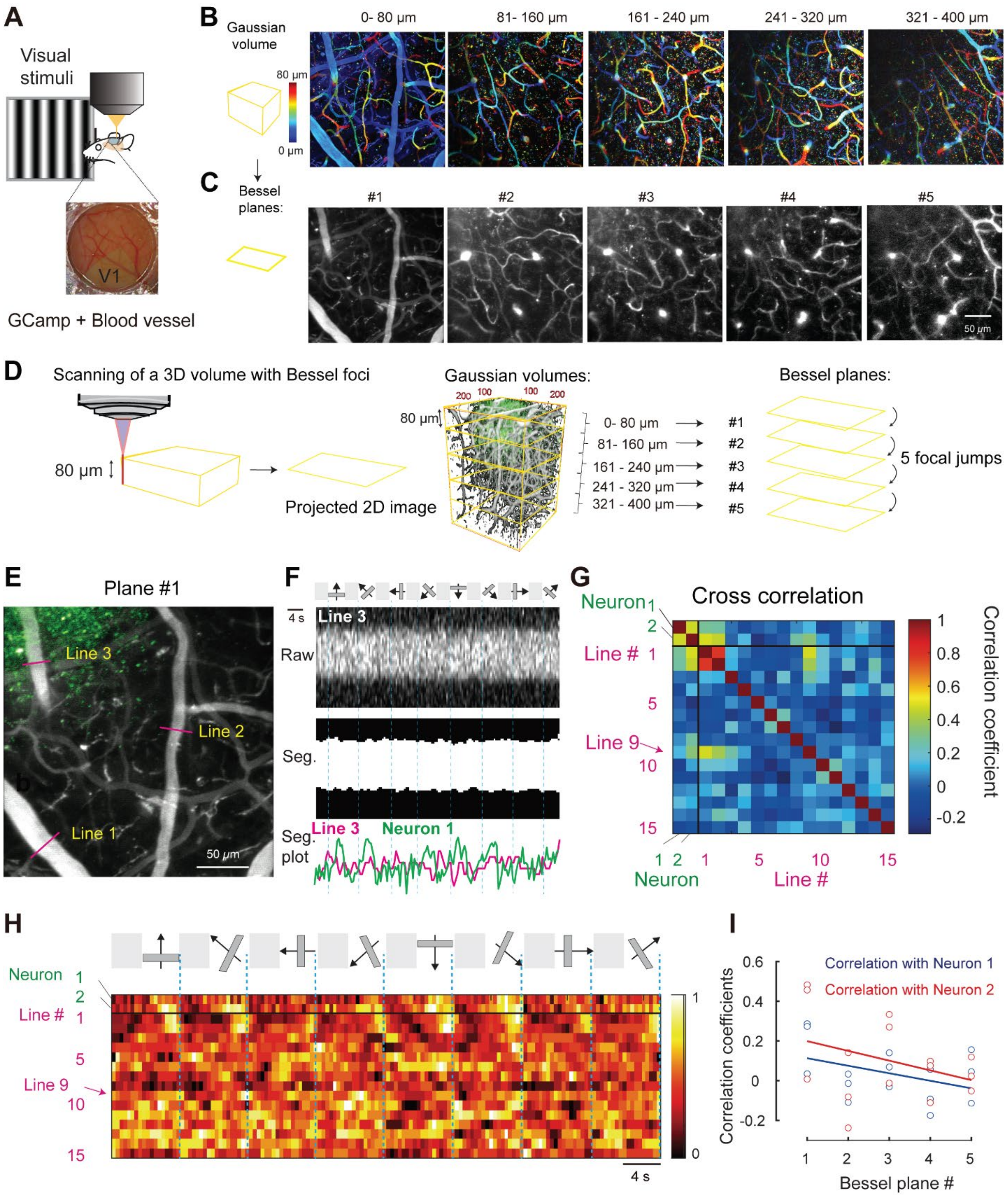


**Figure S7 Neurovascular coupling in response to visual stimulation.** (**A**) Schematic of experimental setup for functional brain imaging of primary visual cortex (V1). Mice with cranial windows installed over V1 area were imaged while being exposed to moving gratings towards eight directions. (**B**) Mean intensity projection of an 80-μm-thick image stack at different depth, which was collected with Gaussian focus scanning at 2-μm z steps, with structures color-coded by depth. (**C**) Images of the same volume of brain collected by scanning a Bessel focus with the length of 80-μm for each depth. (**D**) Schematic diagram of how to use tBessel foci for cross-layer deep brain imaging. A Bessel focus with the axial length of 80-μm can cover 80-μm deep brain volume. When it is moved by piezoelectric actuators along z direction four times with each time jumping 80-μm, we can cover the complete 0 through 400-μm deep brain volume with 5 Bessel planes at fast speed (2.5-Hz in this case). (**E**) Field-of-view (FOV) from Plane #1 in **Figure 2D** showing where lines were drawn. (**F**) The intensity over Line 3 from a venous across 5 trials aligned by 8 drifting grating angles from 0° to 315°. Raw and segmented images from resliced stacks for Line 3 were shown. Summed intensity plot of lines (Line 3, in magenta) was overlaid with calcium trace of Neuron 1 with correlation coefficient being 0.035. (**G**) Cross-correlation matrix among the two selected neurons and the 15 blood vessels. (**H**) Raster plot of the normalized activities of the two selected neurons and diameter changes of the 15 blood vessels (Line 1-15) in (**D**). (**I**) Correlation coefficients between the temporal curves from the 15 blood vessels and Neuron 1 (blue) or Neuron 2 (red).

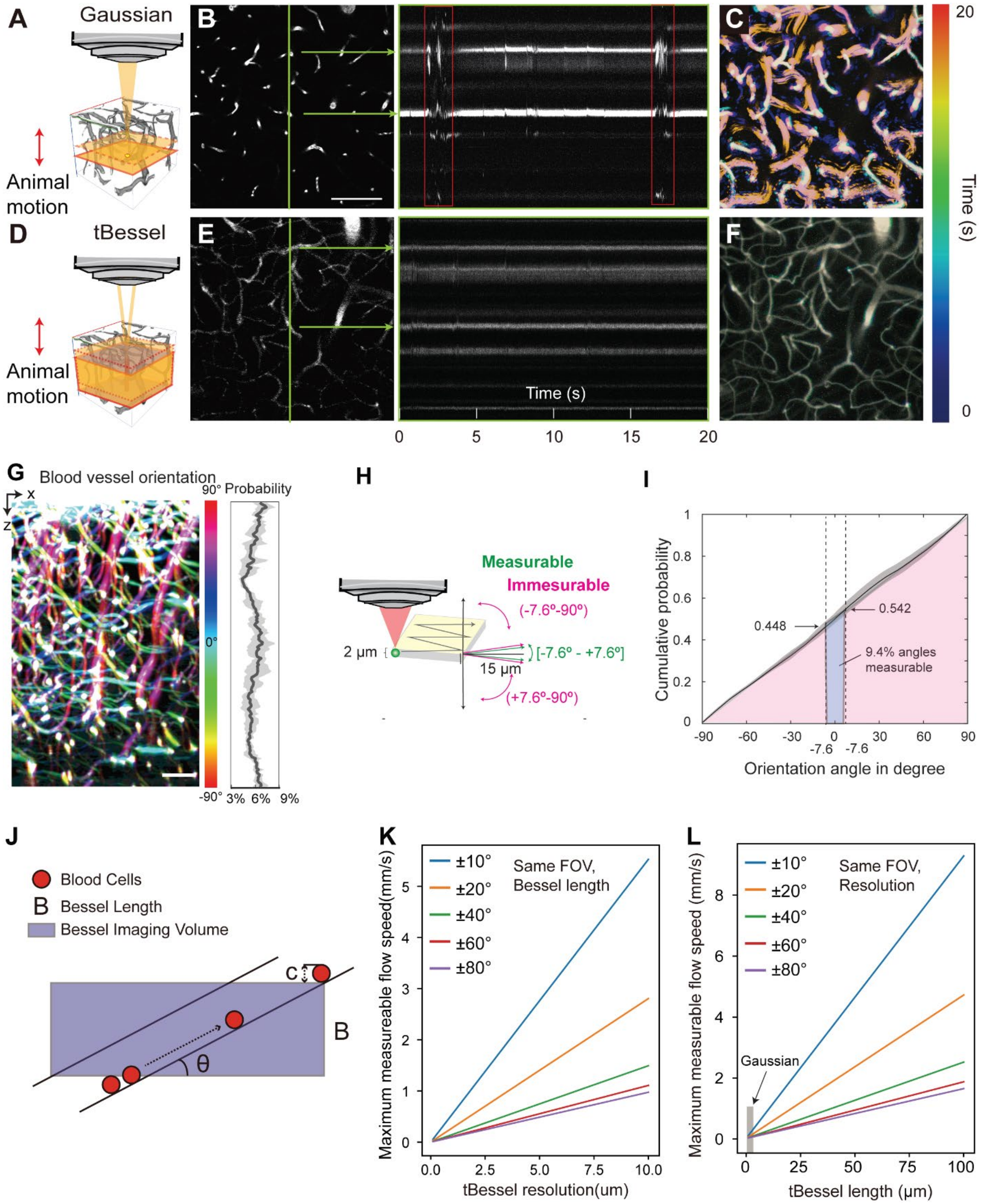


**Figure S8 Tracking hemodynamics in live mouse brains.** (**A**) Schematics of Gaussian imaging: axial motion can move the structures of interest in and out of the focal plane. (**B**) A representative time-lapse vasculature image (400×400-μm$^2$) collected with Gaussian TPFM. Scale Bar: 100-μm. On the right, kymograph of the green line cut. Red boxes highlighted axial motion induced artifacts. (**C**) A color-coded projection of the time-lapse image series. White color indicates structures that stay unchanged, while other colors highlight structural changes due to mouse axial motion. (**D**)-(**F**)**,** Schematics (**D**), representative time-lapse and kymograph (**E**), and color-coded projection (**F**) of the same blood vessel images collected by an 80-μm long tBessel beam. (**G**) Color-coded blood vessel orientation map showing a relatively uniform distribution of blood vessels orientations. Scale bar: 50-μm. (**H**) Diagram of the line-scan measurement under the conventional Gaussian foci with axial point-spread function of 2- μm. When the length of the line at the xy plane is 15-μm, since arctan(2/15) is 7.6$^o$, blood vessels tiling higher than this angle (-7.6 to -90 degrees and -7.6 to -90 degrees) will not be available for line-scan blood flow measurement. (**I**) Cumulative distribution of orientation angles of blood vessels shown in (**G**) averaged over 5 projections. Gray area shows the standard deviation. Blue area indicates the measurable area and red area indicates the not-measurable area in the setting shown in (**H**). Numbers shows the probability correspond to -7.6 and 7.6 degrees at the averaged line. (**J**) Schematic of red blood cells moving through blood vessel with an angle to the horizontal plane. The same red blood cell must be captured between two frames to be registered for blood flow speed measurements. (**K**) Simulated maximum measurable flow speed plotted against tBessel resolution with same FOV and Bessel length, of blood vessels with different orientation angles (color-coded). (**L**) Simulated maximum measurable flow speed plotted against tBessel length of blood vessels with different orientation angles (color-coded). A Gaussian focus is equivalent to a 2-μm long tBessel beam as shown in the grey shaded area.

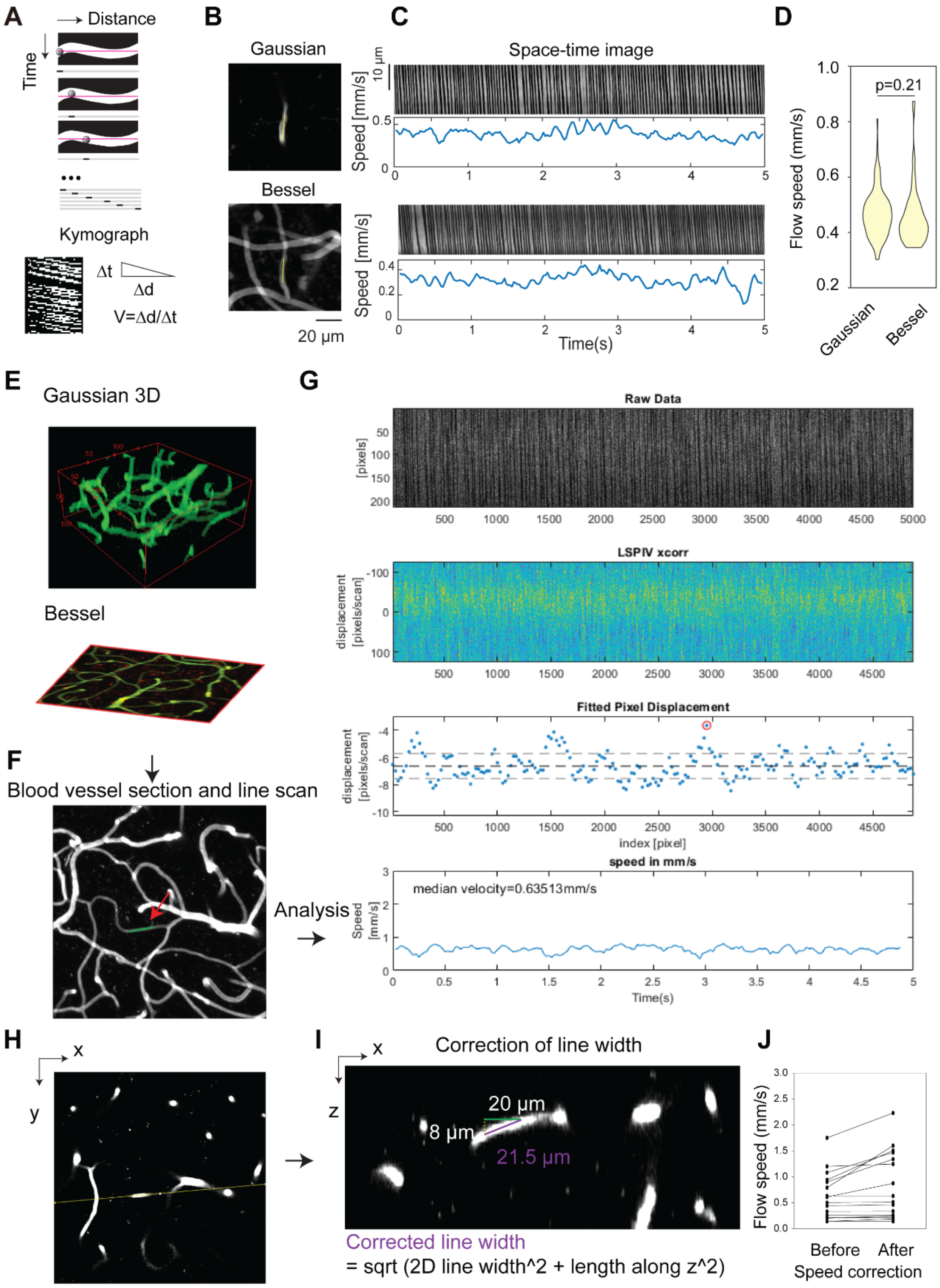


**Figure S9 Measuring blood flow speed with line-scan tBessel-TPFM.** (**A**) Schematic of line-scan for blood flow measurement. The line-scan path is represented as a red line and a stack of sequential line-scans showing the red blood cell at three different locations in the vessel. (**B**) The

2D imaging plane before line-scanning the same blood vessel under Gaussian foci (upper panel) or 80-μm-length Bessel foci (lower panel). (**C**) Space-time images and the plot of blood flow speed as a function of time. The repetition rate of the line-scan is 1 kHz. (**D**) Violin plot showing the distribution of flow speed (in millimeter per second, mm/s) calculated through the streak angle of RBCs in each imaging paradigm. N = 234 angles for Gaussian and 97 for Bessel. P values were calculated by the two-sided Student's t test. $P = 0.21$. (**E**) Gaussian image stack of image of a $200 \times 200 \times 80\ \mu m^3$ volume of vasculature and tBessel-TPFM volumetric imaging of the same volume. (**F**) A segment of blood vessel was chosen for line scan at 1 kHz for 5 seconds. (**G**) Kymographs were generated and analyzed using the line-scanning particle image velocimetry method. Median velocity was calculated. (**H**) and (**I**), the line width was corrected by reslicing the stack and found the spanning range of the line along z direction. The true line width was calculated for correction of the flow speed. (**J**) Blood flow speed before and after correction from 16 line measurements.

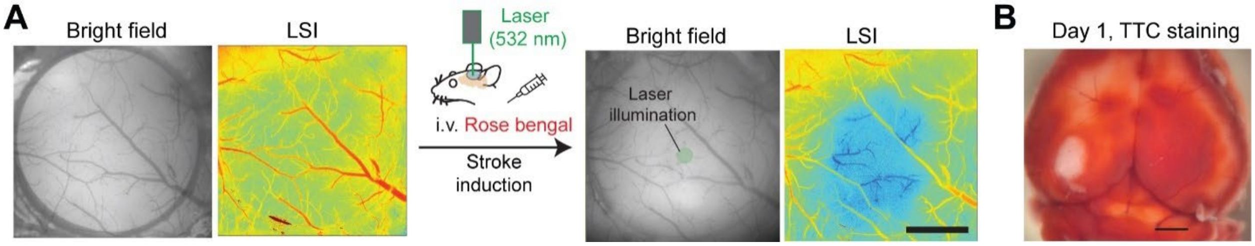


**Figure S10 Characterization of photothrombosis model of ischemic stroke in the mouse visual cortex.** (**A**) The bright field images of the cranial window (3-mm diameter) and LSI images before and after stroke induction. (**B**) TTC staining of the mouse brain 24 hours (day 1) after stroke induction.

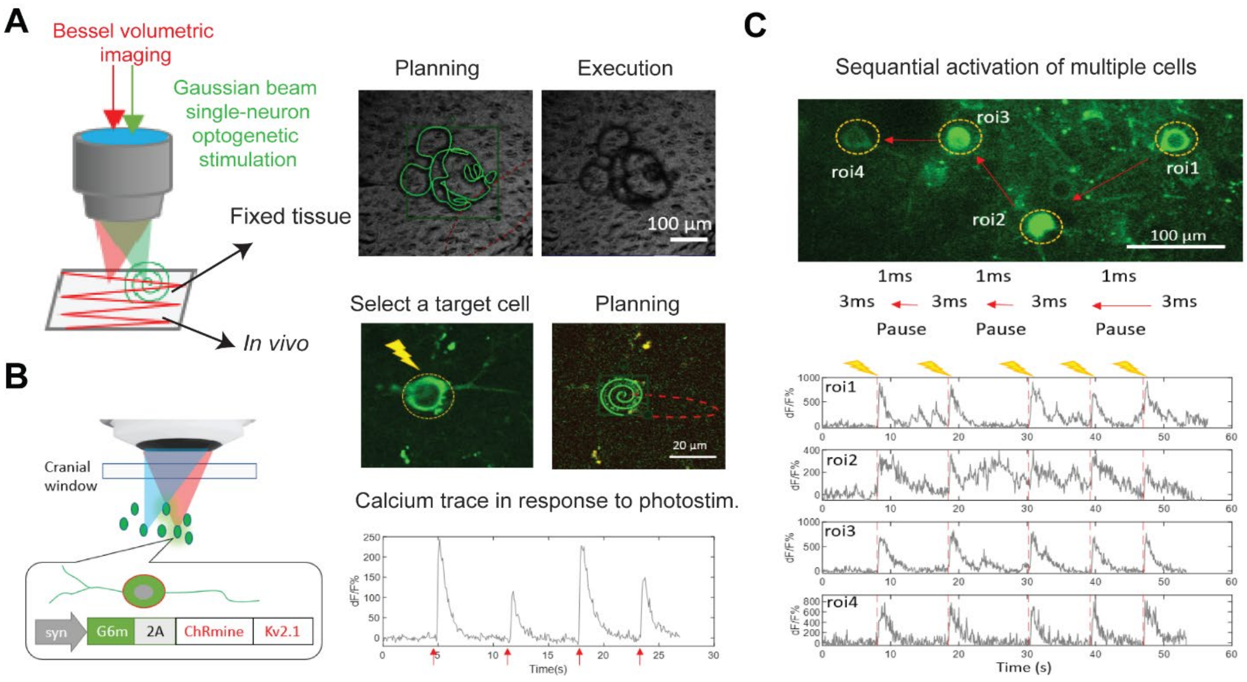


**Figure S11 Integrated optogenetic stimulation and volumetric imaging.** (**A**) Schematic (left) of the experimental configuration. Right: In fixed tissue, arbitrary stimulation patterns can be planned and executed with high precision using the stimulation beam (top). *In vivo* stimulation begins with selecting a target neuron, followed by planning a spiral (vortex) pattern to deliver light precisely to the soma for single-cell activation. (**B**) Schematic of the *in vivo* experimental. Mice expressed a bicistronic construct (syn-GCaMP6s-P2A-ChRmine-Kv2.1) that enables co-expression of the calcium indicator GCaMP6s and the optogenetic actuator ChRmine in the same neurons. Right: Example calcium trace showing reliable responses to repeated optogenetic stimulation (red arrows). (**C**) Sequential stimulation of four individual neurons (3-ms per site with 1-ms inter-site pause). Bottom: Calcium traces from each ROI. Yellow arrows mark stimulation onset. Scale bars: 100-µm.

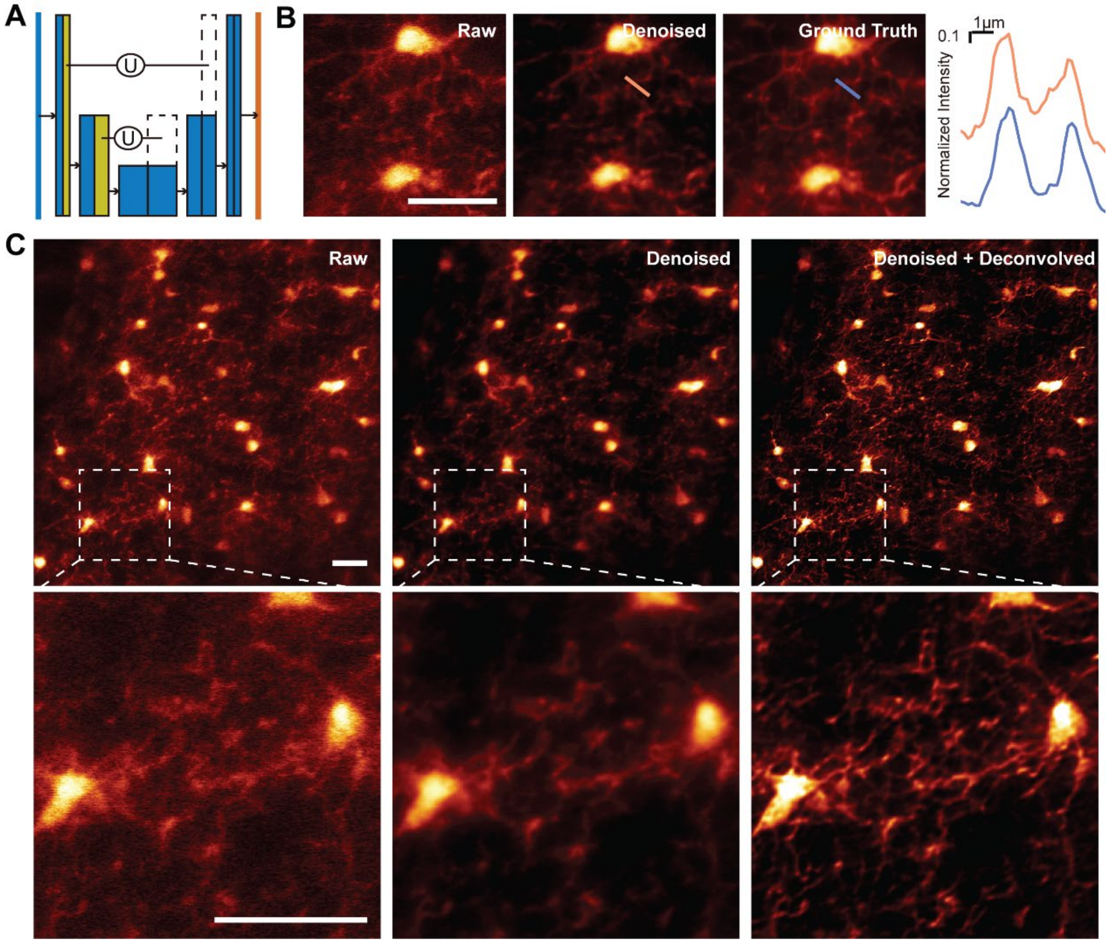


**Figure S12 Denoising and deconvolution pipeline.** (**A**) Schematic of the Content-Aware Image Restoration (CARE)-based U-Net architecture for denoising. Training data were generated by averaging 10 consecutive raw frames to produce high-SNR "ground truth" images, paired with corresponding single-frame low-SNR inputs. (**B**) Representative example comparing raw, denoised, and ground truth images. Line profile analysis (right) demonstrates high concordance with the ground truth. Scale bar: 20-µm. (**C**) Comparison images of raw, denoised, and deconvolved images. Scale bars: 20-µm.

**Table S1. Imaging conditions.**

| Figures | Bessel NA | Bessel length (μm) | Pixel size (μm) | Imaged volume (x,y,z) (μm) | Volume rate (Hz) | Objective | Post-objective power (mW) |
|---|---|---|---|---|---|---|---|
| **2A** | 0.6 | 17-150 | 0.7 | 350,350,500 | *NA* | 16× | 65 |
| **2B,C** | 0.66 | 150 | 1.3 | 2500,2500,450 | 3 (each subvolume) | 16× | 20-92 |
| **2D** | 0.4 | 80 | 0.5 | 260,260,400 | 2.5 | 25× | 47-149 |
| **3B** | 0.4 | 80 | 0.4 | 100,80,80 | *NA* | 25× | 47 |
| **3C,D** | 0.4, 0.6, 0.8 | 60-120 | 1.5 | 400,400,120 | 58，44，29 | 25× | 47 |
| **3G,J,K** | 0.4 | 140 | 0.26 | 1,400,1,400,140 | 1,000 (line-scan) | 16× | 72-98 |
| **5D-F** | 0.7 | 120 | 0.19 | 200,200,120 | 15 | 25× | 70 |
| **S7B,E** | Same as 2D | | | | | | |
| **S8E,F** | 0.66 | 80 | 1.3 | 400,400,80 | 30 | 25× | 47 |
| **S9B** | 0.4 | 80 | 0.3 | 60,60,80 | 1,000 (line-scan) | 25× | 23 |
| **S9F,H** | 0.4 | 80 | 0.4 | 200,200,80 | 1,000 (line-scan) | 25× | 47 |
| **S12B,C** | Same as 5D | | | | | | |
| **MovieS2** | 0.4 | 150 | 1.4 | 1400,1400,150 | 15 (spontaneous), 3(visual stimulation) | 16× | 32 |
| **MovieS3** | Same as Figure S8E,F | | | | | | |
| **MovieS4** | Same as Figure 3D | | | | | | |
| **MovieS5** | Same as Figure 5D | | | | | | |

**Movie S1.** Independent continuous adjustment of NA and ΔNA with the tunable Bessel module.

**Movie S2.** Time-lapse imaging of blood vessel for blood flow velocity measurement over a 400×400×120-μm$^3$ volume at 58-Hz.

**Movie S3.** Time-lapse hemodynamics imaging over a 1,400×1,400×150-μm$^3$ volume showing spontaneous neurovascular activities at 15-Hz and responses to moving grating stimulations at 3-Hz.

**Movie S4.** Time-lapse imaging showing direct comparison of axial motion artifact between Gaussian-TPFM (left) and Bessel-TPFM (right) over a 400×400-μm$^2$ field of view at 30-Hz.

**Movie S5.** Time-lapse imaging of microglia responses immediately after microablation of one microglial cell over a 200×200×120-μm$^3$ volume at 15-Hz for 10-minutes.
